# Genomic profiling of active vitamin D colonic responses in African- and European-Americans identifies an ancestry-related regulatory variant of *POLB*

**DOI:** 10.1101/2025.04.09.647365

**Authors:** David Witonsky, Bharathi Laxman, Hina Usman, Margaret C. Bielski, Kristi M. Lawrence, Sonia S. Kupfer

## Abstract

We measured genomic responses to active vitamin D, 1α,25-dihydroxyvitamin D (1,25D), in colonic organoids from individuals of African and European ancestry. Given protective effects of 1,25D for gastrointestinal conditions such as colorectal cancer, organoid cultures enabled evaluation of condition-specific responses in relevant target tissue. We found significant alterations in transcriptional and chromatin accessibility responses to 1,25D treatment, including some with ancestry-associated differences, and also elucidated the role of *cis-*genetic variance on treatment responses. Integration of genomic profiling with genetic mapping found an insertion-deletion variant that explains ancestry-associated differences in 1,25D regulation of *POLB*, an oxidative DNA repair enzyme involved in colorectal carcinogenesis, which also showed signals of positive natural selection. These findings highlight the importance of including diverse individuals in functional genomics studies to identify potential drivers of population-level differences relevant for clinical outcomes, and to uncover functional mechanisms that may be obscured by ancestry variation.

## Introduction

Active vitamin D, known as calcitriol or 1α,25-dihydroxyvitamin D (1α,25(OH)_2_D_3_ or 1,25D), is a nuclear hormone with effects on a variety of biological processes^1^, including protective effects against gastrointestinal (GI) conditions such as colorectal cancer^2-4^, inflammatory bowel disease^5^, and gut dysbiosis^6^. Circulating levels of the inactive form of vitamin D, 25-hydroxyvitamin D (25(OH)D or 25D), vary substantially across ancestries^7^ and may partially account for inter-ethnic variation in the prevalence and severity of multiple vitamin D-linked disorders. However, while serum 25D levels are a proxy for overall vitamin D stores, an exclusive focus on this measure may obscure clinically significant aspects of the active vitamin’s impact on human biology. We propose that both individuals and ancestry groups may display substantial variation in the responses of target tissues to 1,25D, irrespective of circulating levels. An individual’s genotype and ancestral background could be an important determinant of their response to 1,25D which could influence clinical outcomes and might explain the ambiguous results of vitamin D supplementation trials that fail to control for ancestry and other sources of inherited variation^8-10^. Genomic measures of individual responsiveness to 1,25D, such as differences in chromatin accessibility and transcriptional activity, may therefore be more relevant clinical endpoints than 25D levels^11^.

Prior efforts to characterize genomic responses to vitamin D in humans have mainly been conducted in systems that are relatively easy to sample, such as peripheral blood cells^12,13^ and cell lines^14-16^. While these investigations may yield meaningful results for certain cell types, they may not recapitulate the context-specific responses in relevant tissues across multiple individuals, limiting their applicability to assess inter-individual responses. Alternative models in relevant tissue types from genetically diverse individuals may enable improved characterization of responses, as we demonstrated in a previous study in primary colon organ cultures^17^.

However, those cultures were limited by short viability and mixtures of epithelial and stromal cells. Organoid cultures, derived from colonic epithelial stem cells with longer and more stable viability^18^, offer a more robust experimental framework for studying genomic responses in human colon to environmental factors including 1,25D^19^. Observations of coordinated shifts in chromatin accessibility and transcriptional activity in these cultures following experimental perturbations can be particularly helpful for revealing specific regulatory pathways controlled by treatments of interest such as vitamin D across individuals.

Here, we applied this approach to the question of inter-individual and inter-ancestry variation in 1,25D responses within the colon, comparing treatment responses in colonic organoid cultures from individuals of African and European ancestry. By comprehensively profiling transcriptional and chromatin accessibility responses to 1,25D across heterogenous individuals, we identified novel context-specific genetic effects that regulate these molecular responses, including ancestry-related regulation of biologically relevant genes for GI health and disease.

## Results

### Vitamin D treatment induces widespread genomic responses in colonic organoids

We characterized genomic responses to 1,25D by comparing gene expression and chromatin accessibility data from vitamin D- and control-treated organoids (**Figure 1a**). Colonic biopsies from 60 healthy patients were used to generate colonic organoids. After 24 hours in differentiation media, we separately treated replicates of each of the 60 organoid lines with either 100nM 1,25D or vehicle control (0.1% ethanol) based on the results of our pilot study where we had determined optimal treatment dose and duration for this system^16^. We prepared organoids for ATAC-seq and RNA-seq after 4 and 6 hours of exposure to each treatment, respectively. For transcriptional responses, after quality controls (see Methods), we retained a total of 53 lines for transcriptome profiling, including 26 African-Americans (AA) and 27 European-Americans (EA). For chromatin accessibility profiling, we applied additional stringent quality controls (see Methods) and retained ATAC-seq data from 25 individuals, including 12 AA and 13 EA donors (**Supplementary Table 1**).

**Figure 1:**
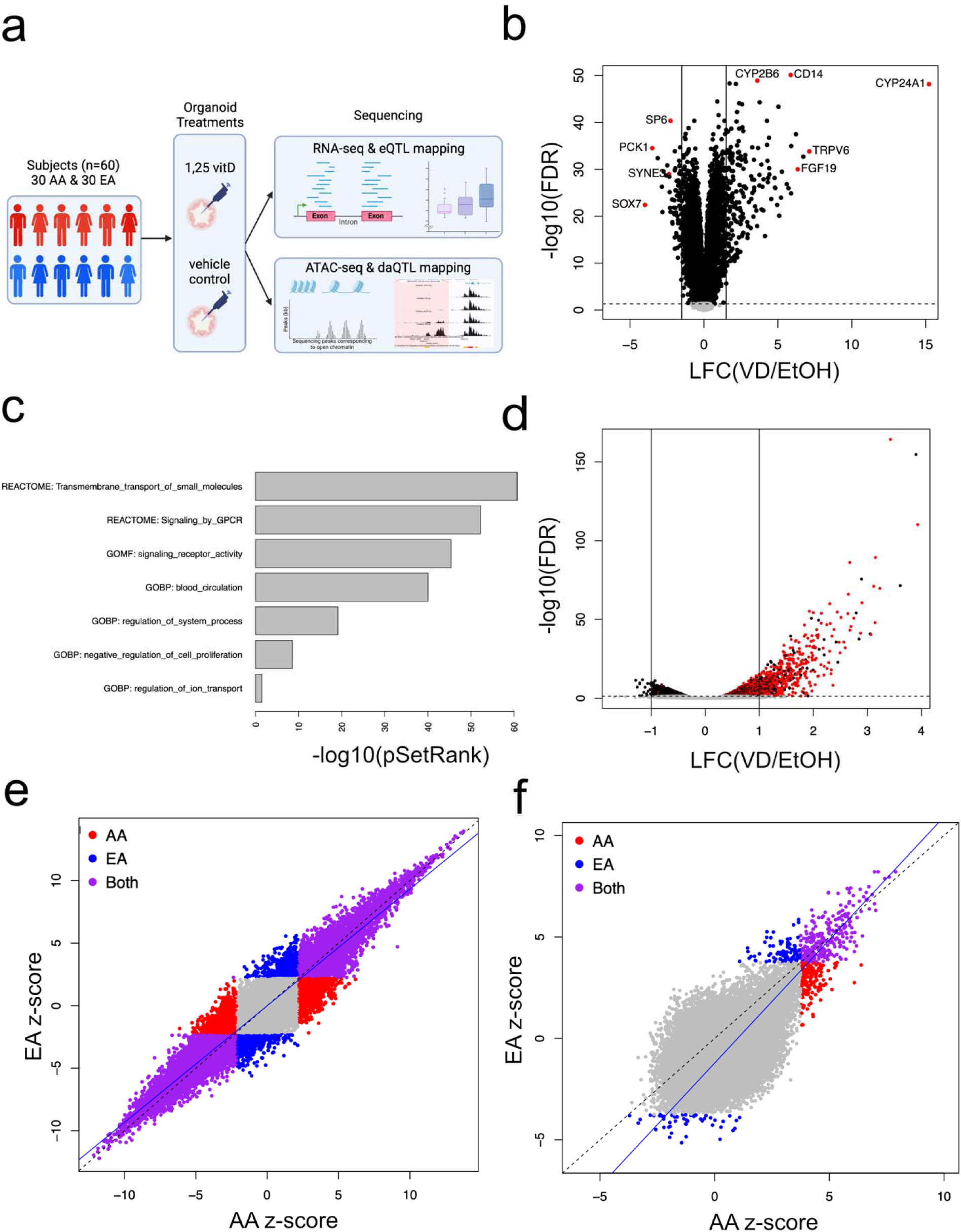
Genomic responses to 1,25D treatment in colonic organoids. **1a) Study design:** Colonic biopsies from a total of 60 healthy patients were obtained during colonoscopy screening exams from which colonic organoids were generated. After 24 hours in differentiation media, we cultured replicates of each organoid line with either 100nM 1,25D or vehicle control (0.1% ethanol). After 4 and 6 hours of exposure to each treatment, we performed ATAC-seq and RNA-seq, respectively. We then mapped QTLs for condition-specific responses (i.e., eQTL and daQTL). Figure created in biorender. **1b) Differentially expressed (DE) genes with 1,25D treatment.** Of 13,163 protein-coding genes tested, 9,066 DR genes were identified at FDR<5% (black dots). Among genes with larger effect sizes (e.g., |LFC|>1.5; indicated by horizontal lines), there was a 3-fold greater upregulation compared to downregulation. Top upregulated genes included known targets of the VDR such as *CYP24A1, TRPV6, FGF19, CYP2B6*, and *CD14*, while top down-regulated genes included *SOX7, PCK1, SYNE3*, and *SP6* (red dots). **1c) Gene set enrichment analysis of transcriptional responses.** A total of 128 pathways from 3 databases (KEGG, GOBP, REACTOME) were significantly enriched among DE genes based on adjusted p-values. Significantly enriched pathways included functions related to transmembrane transport, signaling by G protein coupled receptors as well as negative regulation of cell proliferation and regulation of ion transport. **1d) Differential chromatin accessibility (DA) with 1,25D treatment.** A total of 4,142 peaks (3.5%) were DA with 1,25D treatment at FDR<5% (black dots). Of 639 strong DA peaks (e.g., |LFC| >1) (indicated by horizontal lines), 622 (97.5%) showed greater accessibility with 1,25D treatment. The motif for the vitamin D response element (VDRE, defined by a predicted VDR and RXR binding; red dots) was significantly enriched among DA peaks, especially those with positive LFC. **1e) Genome-wide 1,25D transcriptional responses by ancestry.** There was significant correlation between z-scores for DE genes with treatment between AA and EA (r=0.96; p<2.2x10^-16^). A large number of DE genes were significant in both populations at FDR<5% (purple dots). A smaller number of DE genes were significant in AA (red dots) or EA (blue dots). Line y = x (dashed line). Line of best fit for DE genes: y = -0.01 + 0.93 (blue line). **1f) Genome-wide 1,25D chromatin accessibility by ancestry.** Similar to findings for DE genes, we found strong correlation between z-scores for DA peaks (r=0.86; p<2.2x10^-16^). When testing for DA peaks in the populations individually, a total of 582 DA peaks were found in AA or EA at FDR<5%. Of these, 177 DA peaks were significant in AA only (red dots), 141 DA peaks in EA only (blue dots) and 264 DA peaks were significant in both populations (purple dots). One peak located proximal to the *POLB* promoter (circled dot) showed a significant difference in DA LFC between populations. Line y = x (dashed line). Line of best fit for DA peaks: y = -1.21 + 1.21 (blue line).

To identify differentially expressed (DE) genes between 1,25D and vehicle control treatments, we used linear mixed models implemented in the *dream* R software package^20^, including individual as a random effects variable and age, sex, batch, ancestry, and cell type composition as fixed effects covariates (see Methods Model 1). *FABP1*, a highly expressed enterocyte marker, was used to control for cell type (**Extended data Figure 1a**). All organoid lines displayed substantial shifts in expression with 1,25D exposure (**Extended data Figure 1b**). Of 13,163 protein-coding genes tested, 9,066 DE genes were identified at a false discovery rate (FDR) <5% (**Figure 1b**). Of these, 254 (2.8%) showed an absolute log2 fold change (LFC) with magnitude 1.5 or greater. This highly DE subgroup showed greater upregulation than downregulation, with 187 genes showing increased expression and 67 showing diminished expression — an approximately 3-fold difference. The top upregulated DE genes included genes that are known vitamin D target genes via VDR binding: *CYP24A1, TRPV6, FGF19, CYP2B6*, and *CD14*. Top downregulated DE genes included *SOX7, PCK1, SYNE3*, and *SP6* (full list in **Supplementary Table 2**). The gene encoding VDR showed a small but significant upregulation with 1,25D treatment (LFC=0.36; FDR=3.6x10^-11^).

We performed gene set enrichment analysis of transcriptional responses to characterize the types of biological pathways activated or suppressed in response to 1,25D using SetRank^21^. In total, 128 pathways from 3 databases (KEGG, REACTOME, GOBP) are significantly enriched among DE genes based on adjusted p-values (**Supplementary Table 3**). Top enriched pathways include functions related to transport, signaling by G protein coupled receptor as well as negative regulation of cell proliferation and ion transport (**Figure 1c**). One disease-associated pathway was significantly enriched among DE genes, namely “pathways in cancer” (KEGG hsa05200) (**Extended data Figure 1d**).

Next, we examined differences in chromatin accessibility between treatment and control conditions across 118,806 ATAC peaks detected in at least 3 of the 50 samples from both treatment conditions. We identified differentially accessible (DA) peaks at an FDR<5% using a linear model implemented in the DiffBind R package^22^, including individual as a fixed effect covariate (see Methods Model 3). Similar to gene expression, organoid lines displayed substantial shifts in chromatin accessibility with 1,25D exposure (**Extended data Figure 1c**). We found 4,142 DA peaks (3.5%), of which 639 (15.3%) displayed high responsiveness (|LFC| >1). The vast majority of this subset of highly responsive DA peaks (622 peaks, 97.5%) showed greater accessibility in response to 1,25D (**Figure 1d; Supplementary Table 4**). DA peaks showed significant enrichment relative to all peaks in intronic (49.3% vs. 43.8%, respectively; p=2.10x10^-13^) and intergenic (41.7% vs. 38.0%, respectively; p=3.32x10^-7^) regions. Conversely, DA peaks were significantly depleted in promoter regions (3.6% vs. 12.4%, respectively; p=1.54x10^-89^). This depletion was also seen reflected in a broader distribution of distances to the transcription start site (TSS) of the nearest gene for DA peaks relative to peaks that were not DA (**Supplementary Table 5; Extended data Figure 2a**). A total of 85 (0.65%) of the tested protein-coding genes were found to have DA peaks falling in their promoters. Of these, 73/85 (86%) were DE, a 1.3-fold enrichment over genes without DA peaks in their promoters (p=8.18x10^-5^). We found that the corresponding DE and DA effect sizes for these 73 genes were strongly correlated (r=0.85; p<2.2x10^-16^) (**Extended data Figure 2b**), suggesting a probable mechanistic link between 1,25D-induced alterations in gene expression and proximal chromatin accessibility. Roughly the same fraction of DE genes with a DA promoter showed upregulation (48%) as the overall set of DE genes. Despite the observed enrichment of upregulated genes among highly DE genes, this more proximal mediation in itself does not alter the likelihood of the direction of gene regulation.

**Figure 2:**
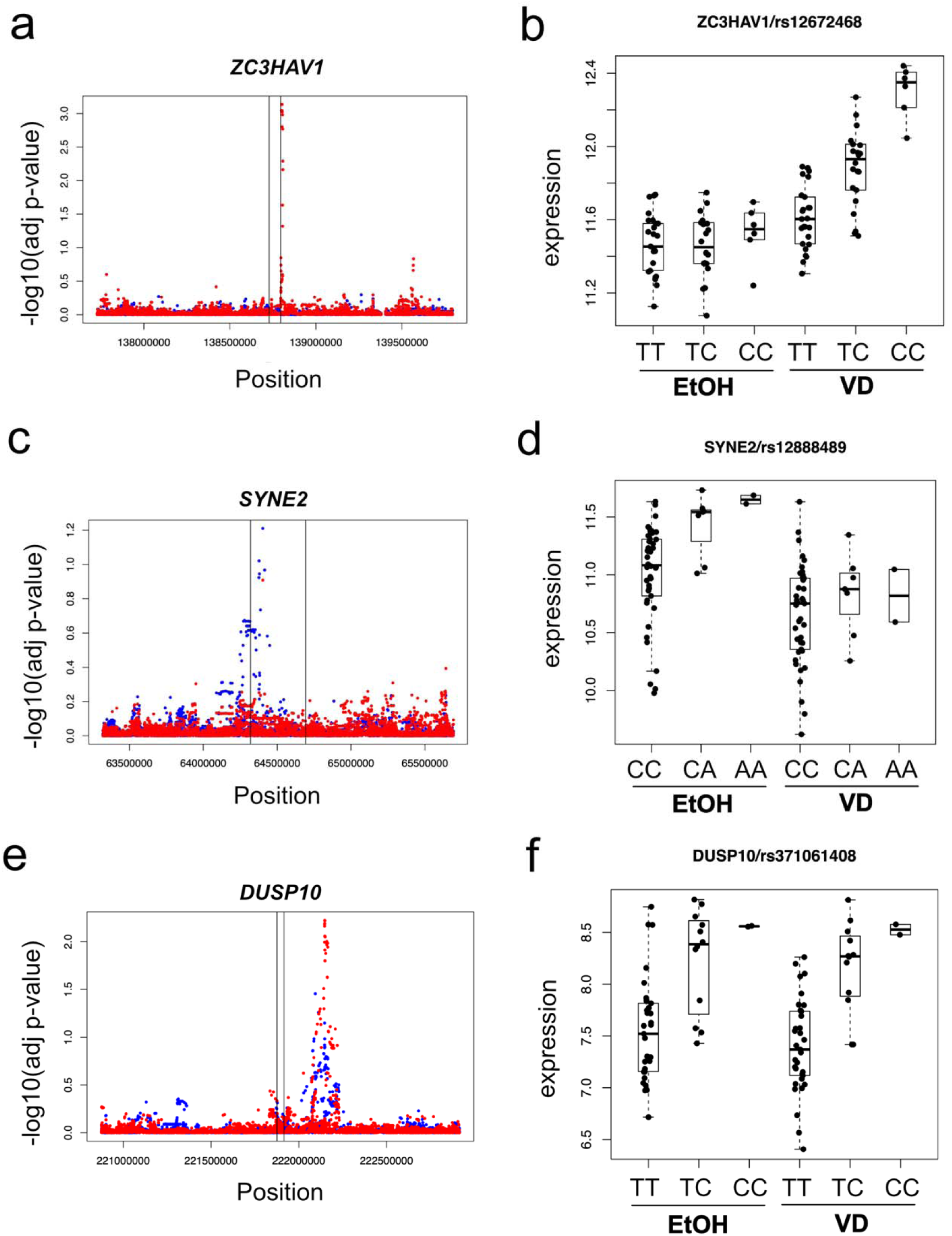
*cis-*eQTL mapping of condition-specific 1,25D treatment responses. *cis-*eQTL mapping of condition-specific treatment responses was performed using BRIdGE, a Bayesian approach designed to assess gene-environment interactions for paired measurements under two conditions. Mapping was done using genotyped/imputed SNPs. At a posterior probability cut-off of >0.70, BRIdGE identified 132 vitamin D-specific eQTLs and 41 control-specific eQTLs. A total of 2522 *cis-*eQTLs were significant in both conditions. Examples of condition-specific eQTLs are shown in this figure. MatrixEQTL adjusted p-values were plotted as a function of physical position for 1,25D (red dots) or control (blue dots) treatment conditions. Box plot expression is the DESeq2 variance stabilizing transformation (vst) of normalized read counts. **2a & b) Example of 1,25D-specific *cis-*eQTL**. A number of significant eQTLs on chromosome 7 were found only with 1,25D treatment (**2a**). Of these, rs12672468 showed the highest posterior probability for vitamin D-specific responses for *ZC3HAV1*, a zinc-finger protein located that has been implicated in colorectal cancer and innate immunity. A genotypic effect was noted for 1,25D-treated samples but not for control treatment (**2b**). **2c & d) Example of control-specific *cis-*eQTL**. On chromosome 14, an enrichment was noted for significant eQTLs predominantly with control treatment (**2c**). Of these, rs12888489 showed the highest posterior probability for control-specific responses for *SYNE2*, a spectrin repeat containing protein. This protein forms a linking network for cellular spatial organization. *SYNE2* was found to be a somatically methylated gene in colon cancer and associated with colon cancer sidedness and prognosis. A genotypic effect was noted only among control-treated samples but not for 1,25D treatment (**2d**). **2e & f) Example of *cis-*eQTLs in both condition**. The largest number of *cis-*eQTLs were associated with expression of genes in both treatment conditions. On chromosome 1, there was enrichment for significant eQTLs in both treatment conditions (**2e**). Of these, rs371061408 was associated with expression of *DUSP10* with 1,25D and control treatment. *DUSP10* encodes a dual-specificity protein phosphatase that negatively regulates MAP kinases that has been significantly associated with colon cancer in previous genetic association studies. For this gene, genotypic effects on expression were seen for both 1,25D- and control-treated samples (**2f**). **Abbreviations:** LFC, log fold change; FDR, false discovery rate; VD, 1,25D vitamin D; EtOH, ethanol vehicle control

We further analyzed the sequence content of the complete set of DA peaks to identify regulatory elements potentially controlled by 1,25D and identified significant enrichment for motifs of 79 transcription factors (TF) using *Homer^23^*. The motif for the vitamin D response element (VDRE), defined by a predicted binding site of VDR and its partner RXR, was present in 19.7% of DA peaks compared to only 2.3% of background peaks (p=1.0x10^-481^; **Supplementary Table 6**).

VDRE was the most significantly enriched motif among DA peaks relative to background peaks with large effects (i.e., LFC>1; 62% vs 2.8%; p=1.0x10^-423^) as well as for peaks with weaker but positive effects (i.e., 0<LFC<1; 30.6% vs 2.5%; p=1.0x10^-313^). The VDRE motif showed a non-significant deficit relative to background in peaks with reduced effects with 1,25D treatment (i.e., LFC<0; 0.74% vs 3.02%; p=1.0). Mirroring these enrichment patterns, a VDRE motif was found within 250 bp of a DA peak center for 75% of DA peaks with large effects, 46% with moderate effects, and only 3.8% of DA peaks with reduced effects (**Extended data Figure 2c**). In addition to VDRE, 39 TF motifs were enriched with 1,25D treatment for DA peaks with large effects (**Table 1**). For peaks with opposite effects with 1,25D, the most significantly enriched motif was for KLF5 (52.3% vs 34.1%; p=1.0x10^-66^). We note that KLF5 expression is significantly downregulated by 1,25D, which could explain the reduced chromatin accessibility under vitamin D treatment for peaks with the KLF5 motif.

**Table 1:**
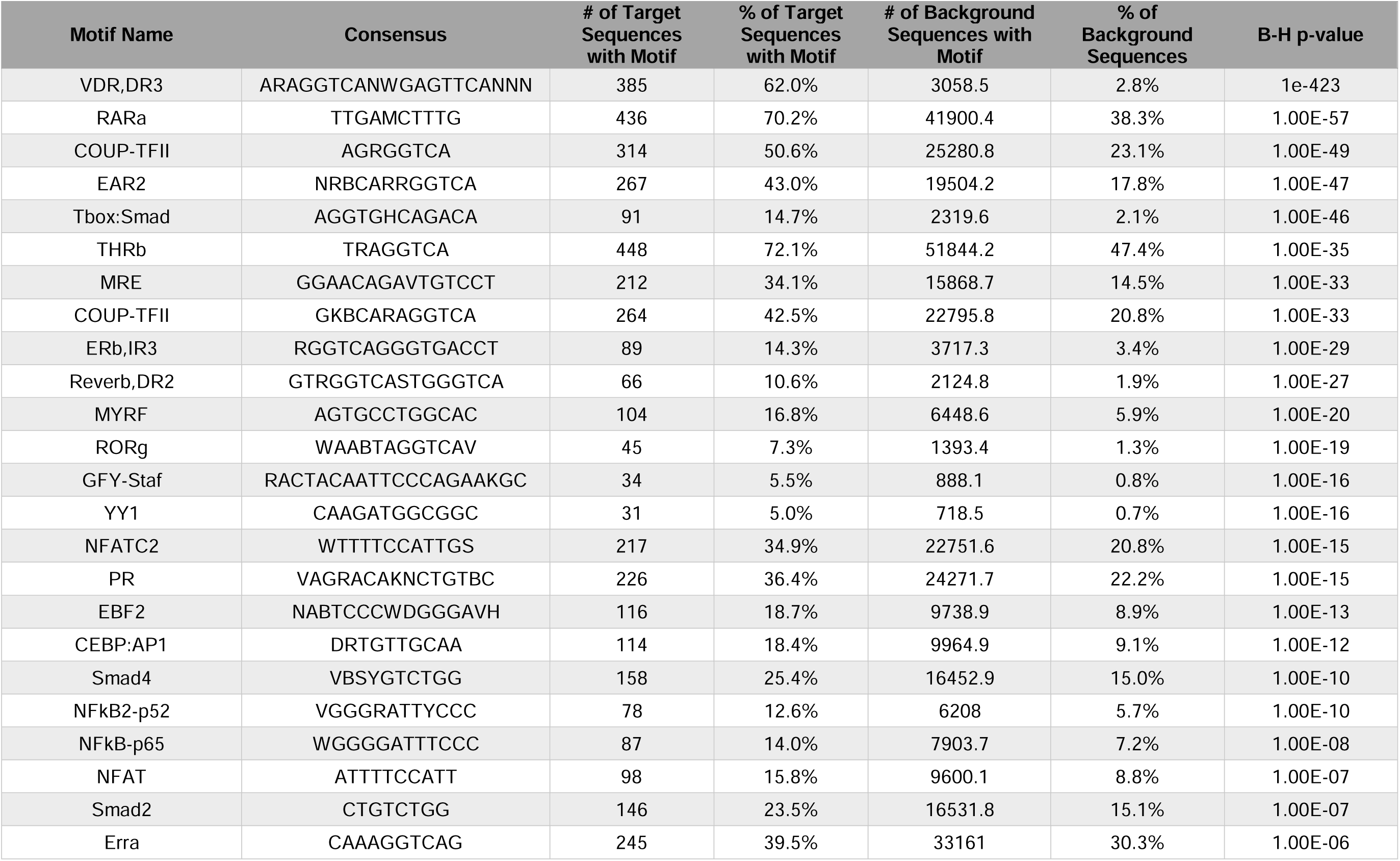
Transcription Factor Motif Enrichment. Top enriched motifs generated from HOMER. Abbreviation: B-H, Benjamini-Hochberg correction for multiple testing

### Inter-ancestry differences in molecular responses to Vitamin D treatment

To test for ancestry-related differences in 1,25D genomic responses, we compared DE and DA results between organoid lines from individuals of African and European ancestry (ancestry proportions shown in **Extended data Figure 3a-b**). We did not observe a large genome-wide difference in the transcriptional responses to 1,25D between AA and EA individuals, as shown by strong correlations both between z-scores (r=0.96; p<2.2 x 10^-16^) (**Figure 1e**) and effect sizes (r=0.97; p<2.2 x 10^-16^) (**Extended data Figure 3c**) across DE genes. Furthermore, neither population showed a stronger overall absolute response to treatment, as demonstrated by a non-significant between population transcriptional effect size magnitude difference (|LFC_AA_| - |LFC_EA_|), when compared to the expected null distribution generated from permuted samples (p=0.28) (**Extended data Figure 3e**).

**Figure 3:**
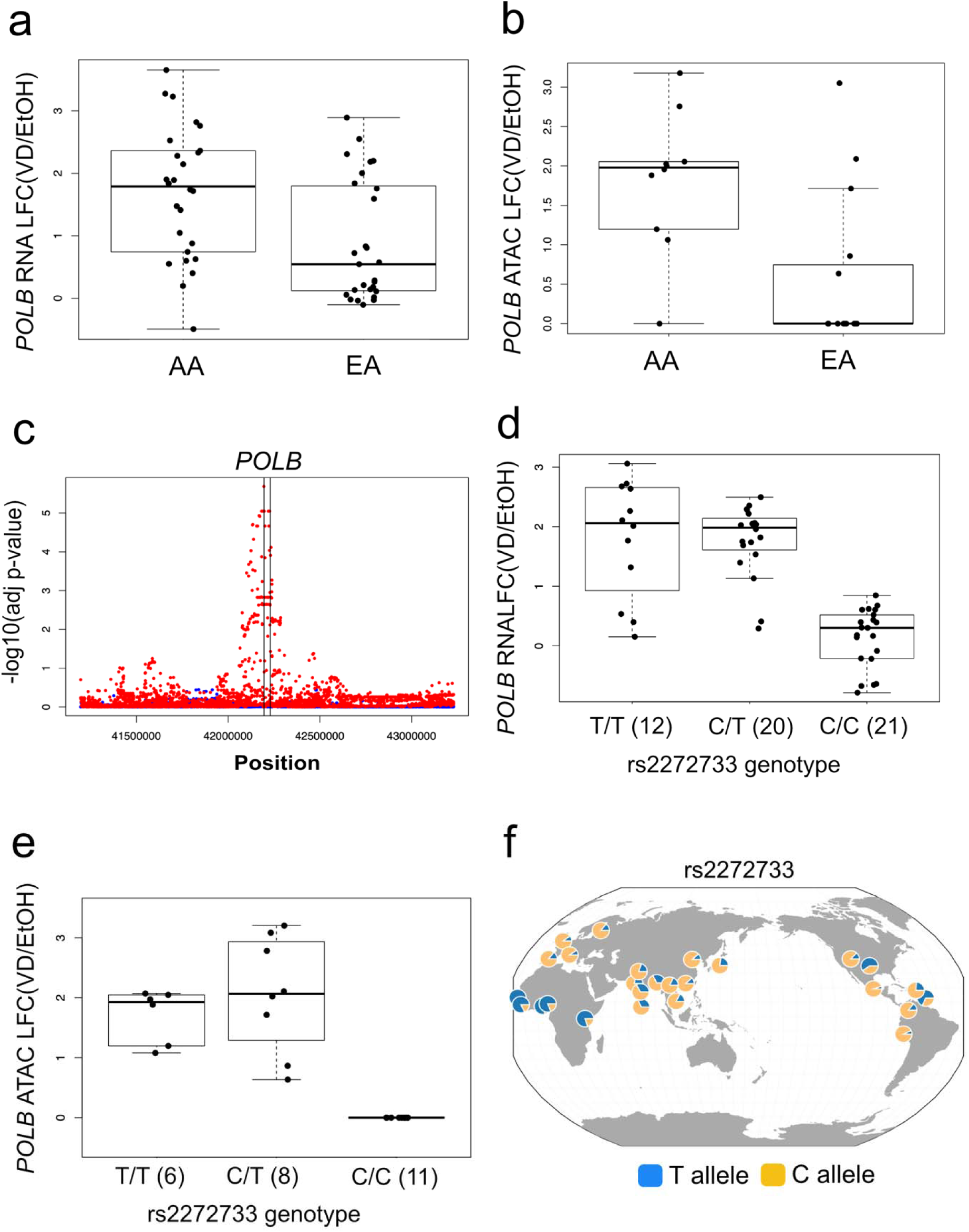
*POLB* vitamin D responses differed by ancestry and showed a significant vitamin D-specific *cis-*eQTL with large allele frequency differences between African and non-African populations. **3a) Differential 1,25D transcriptional responses of *POLB* differ by ancestry.** We noted greater 1,25D transcriptional responses of *POLB* in AA compared with EA lines. In AA lines, the *POLB* LFC for 1,25D response was 1.91 (p=1.12 x 10^-13^), compared to EA lines in which *POLB* LFC was 1.05 (p=3.03 x 10^-6^). Comparison of *POLB* LFC in AA/EA was 0.86 (p=0.003). **3b) Differential 1,25D accessibility peak near *POLB* differs by ancestry.** The most proximal DA peak to the *POLB* gene (“peak A”), lying approximately 2.3 kb upstream of the *POLB* TSS, was noted to have greater accessibility in response to 1,25D in AA (p=2.6 x 10^-7^) compared with EA lines (p=3.8 x 10^-1^). The difference between the two populations was significant (t-test p=3.2 x 10^-3^). **3c) *POLB* vitamin D-specific response *cis-*eQTL.** From our daQTL mapping, we found that only the *POLB* proximal peak had significant daQTLs that were also significant eQTLs for a protein-coding DE gene, namely *POLB*. One of these daQTLs is the significant vitamin D-specific response eQTL rs2272733, a SNP genotyped in our dataset, for *POLB.* MatrixEQTL adjusted p-values were plotted as a function of physical position for 1,25D (red dots) or control (blue dots) treatment conditions. **3d) rs2272733 as a vitamin D-specific eQTL for *POLB****. POLB* shows differential responses to 1,25D by rs2272733 genotype. The LFC for T/T and T/C genotypes were significantly increased with 1,25D treatment, while he LFC for the C/C genotype was not significantly different from 0 (mean LFC=0.141, p=0.21). **3e) rs2272733 as a vitamin D-specific daQTL for ATAC peak located proximal to *POLB*.** The ATAC peaks shows differential chromatin accessibility to 1,25D by rs2272733 genotype. There were 0 peak reads under both treatments for the C/C genotype. **3f) *POLB* eQTL, rs2272733, shows large allele frequency differences in African vs. non-African populations.** The significant *POLB* eQTL, rs2272733, was notable for large allele frequency differences in African vs. non-African populations. Depicted in this panel are allele frequencies in global populations (T allele=blue; C allele=yellow). In African populations, the average T allele frequency is 77.3%, while the C allele frequency is 22.7%. In European populations, the average frequencies for T and C alleles are 12.4% and 87.6%, respectively. Visualized using the Geography of Genetic Variants Browser. **Abbreviations:** LFC, log fold change; EA, European-American; AA, African-American; adj p-val, adjusted p-value; Pos, position; VD, 1,25D vitamin D; EtOH, ethanol vehicle control

In order to identify individual genes with ancestry-related differences in 1,25D treatment responses, we applied a mixed model similar to that used to identify DE genes but with an added treatment by ancestry interaction term. This model yielded highly concordant results regardless of whether ancestry was approximated by a dichotomous race variable or a continuous variable of African ancestry proportion (see Methods Models 2a, 2b), as demonstrated by a strong correlation between interaction term z-scores (**Extended data Figure 3d**). We used a nested approach that focused on the subset of genes that had shown both significant differential expression by 1,25D treatment and by ancestry (n=77 genes). A total of 17/77 (22%) genes showed significant differences in treatment response between AA and EA (**Table 2**). For 15/17 (88%) of these genes, responses were greater in AA compared to EA.

**Table 2:** Ancestry-related differences in 1,25D treatment responses. Using genes that differed by treatment and by ancestry (n=77 genes), a mixed model approach was applied to identify DE genes with significant interactions between treatment and population, controlling for individual, treatment, ancestry, age, sex, batch and cell composition. Abbreviations: adj, adjusted; LFC, log fold change (VD/EtOH)

| Gene Name | LFC<sub>AA</sub>-LFC<sub>EA</sub> | p-value | adj p-value |
| --- | --- | --- | --- |
| *HS6ST1* | 0.453 | 4.18E-05 | 3.22E-03 |
| *KYNU* | 1.799 | 1.92E-04 | 7.39E-03 |
| *AMACR* | -0.280 | 5.59E-04 | 1.28E-02 |
| *S100A14* | 0.171 | 6.66E-04 | 1.28E-02 |
| *CDK2AP2* | 0.255 | 1.04E-03 | 1.60E-02 |
| *MRPL28* | 0.199 | 1.44E-03 | 1.85E-02 |
| *ZNF346* | 0.287 | 1.89E-03 | 2.08E-02 |
| *TMEM14C* | 0.140 | 2.22E-03 | 2.14E-02 |
| *POLB* | 0.922 | 2.74E-03 | 2.34E-02 |
| *PITPNA* | 0.120 | 3.48E-03 | 2.68E-02 |
| *C8orf82* | 0.226 | 4.25E-03 | 2.89E-02 |
| *LDHD* | 0.210 | 4.73E-03 | 2.89E-02 |
| *TMEM9* | 0.188 | 4.88E-03 | 2.89E-02 |
| *D2HGDH* | 0.266 | 5.53E-03 | 3.04E-02 |
| *SRMS* | -0.965 | 6.37E-03 | 3.27E-02 |
| *LYN* | 0.259 | 6.86E-03 | 3.30E-02 |
| *POLR2J3* | 0.243 | 8.58E-03 | 3.88E-02 |

In assessment of ancestry-related differences in chromatin accessibility, we noted results similar to those for transcriptional responses, with a strong correlation between the population z-scores across DA peaks (**Figure 1f**; r=0.86; p<2.2 x 10^-16^). When testing for DA peaks in the populations individually, a total of 582 DA peaks were found in AA or EA at FDR<5%. Of these, 177 DA peaks were significant in AA only, 141 DA peaks in EA only and 264 DA peaks were significant in both populations.

### Molecular responses to Vitamin D treatment are genetically regulated

To identify possible genetic contributions to the variation in 1,25D transcriptional responses, we mapped condition-specific *cis-*expression quantitative trait loci (eQTLs) using the program BRIdGE^24^ to analyze genotyped and imputed SNPs. BRIdGE assesses gene-environment interactions for paired measurements under two conditions that explicitly considers specific interaction models (i.e., vitamin D-specific, control-specific or both conditions), and identifies a single SNP with the maximal association for each gene. At a posterior probability cut-off of >0.70, we identified 132 and 41 condition-specific *cis-*eQTLs for 1,25D and control treatments, respectively. A total of 2,522 *cis-*eQTLs were associated in both treatment conditions **(Supplementary Table 7).**

Examples of top condition-specific *cis-*eQTLs are shown in **Figure 2**. Among vitamin D-specific *cis-*eQTLs, rs12672468 showed the highest posterior probability for the gene *ZC3HAV1* (**Figures 2a & b**), a zinc-finger protein previously implicated in colorectal cancer^25^ and innate immune responses^26^. Among control-specific *cis-*eQTLs, rs12888489 showed the highest posterior probability for *SYNE2* responses (**Figures 2c & d**). *SYNE*, a spectrin repeat containing protein implicated in regulation of cell cycle and regulation, has been previously evaluated as a frequently somatically methylated gene in colon cancer^27-29^ associated with colon cancer sidedness and prognosis^30-32^. The largest number of *cis-*eQTLs were associated with expression of genes in both treatment conditions. Of these, rs371061408, associated with *DUSP10* expression (**Figures 2e & f**) was notable given its role in colorectal carcinogenesis, and significant association with colon cancer in previous genetic association studies^33-36^.

To determine whether chromatin accessibility was influenced by genetic variation, we used matrixEQTL^37^ to test for associations between the LFC of each of the 4,142 DA peaks and all SNPs within 100 kb. We applied both linear and ANOVA models to detect additive and dominant effects, respectively (**Supplementary Table 8**). Significantly associated SNPs (FDR<5%) were classified as differential accessibility QTLs (daQTLs). Of the peaks tested, 62 (1.5%) were found to have at least one daQTL under the linear model and 205 (5.0%) under the ANOVA model with 52 peaks having a daQTL under both models. Approximately half (52.2%) of all DA peaks exhibited reduced chromatin accessibility; however, we noted that these peaks accounted for a significantly higher fraction of daQTLs: 40/62 (65%; two-tailed hypergeometric test p=3.6x10^-2^) under the linear model and 146/205 (71%; two-tailed hypergeometric test p=8.49x10^-9^) under the ANOVA. This skew towards daQTLs associated with peaks showing decreased accessibility could be due to TFs mediating a repressed response being more sensitive to genetic variation or the genomic regions that these peaks occupy being less functionally constrained, though these hypotheses require future validation.

To link DNA sequence variation to the underlying regulatory mechanism contributing to variation in 1,25D transcriptional response, we considered SNPs that were daQTLs as well as eQTLs for DE genes in our data. Under both linear and ANOVA models, a total of 34 SNPs that were identified as daQTLs were also found to be eQTLs. These SNPs were all eQTLs for the protein-coding gene *POLB*, a gene that was also strongly DE in our analysis (LFC=1.52; FDR=5.76x10^-13^). These SNPs were also daQTLs all associated with a single peak, which showed strong positive response under 1,25D treatment (LFC=2.42; FDR=2.50x10^-36^). This peak was located approximately 2.3 kb upstream of *POLB*.

*POLB* was of particular interest because of inter-ethnic differences in 1,25D responses. Specifically, we found that the transcriptional response of *POLB* to 1,25D was significantly greater in AA individuals than in EA individuals (**Figure 3a**). Correspondingly, the identified peak exhibited increased chromatin accessibility in AA relative to EA individuals after 1,25D treatment and was found to be DA in the AA subset only (**Figures 2f and 3b**). We reasoned that these differences could be genetically regulated. Of the 34 SNPs that were daQTLs and *POLB* eQTLs, 10 SNPs were found to define a core haplotype where pairwise SNP r^2^>0.90 in both YRI and CEU 1000 Genome Project populations (**Extended data Figure 4a-b**). Notably, one of these SNPs (rs3136717) had already been shown to lie on a haplotype with large allele frequency differences between Africans and Europeans^38^, and another (rs7462320) was identified by BRIdGE as a vitamin D-only *cis*-eQTL for *POLB* with a posterior prob=0.995, the second highest for any tested gene. We selected rs2272733 as a tagging SNP for this haplotype because this SNP was genotyped in our data. Among homozygotes or heterozygotes for the rs2272733 ancestral allele, *POLB* showed significant responses to 1,25D. Compared to individuals with at least one copy of the ancestral allele, homozygotes for the rs2272733 derived allele showed no significant differential *POLB* response with 1,25D treatment (**Figure 3c & d).** A parallel accessibility response by genotype was seen for the associated DA peak **(Figure 3e)**. rs2272733 showed large allele frequency differences between individuals of African vs non-African ancestries (**Figure 3f**).

**Figure 4:**
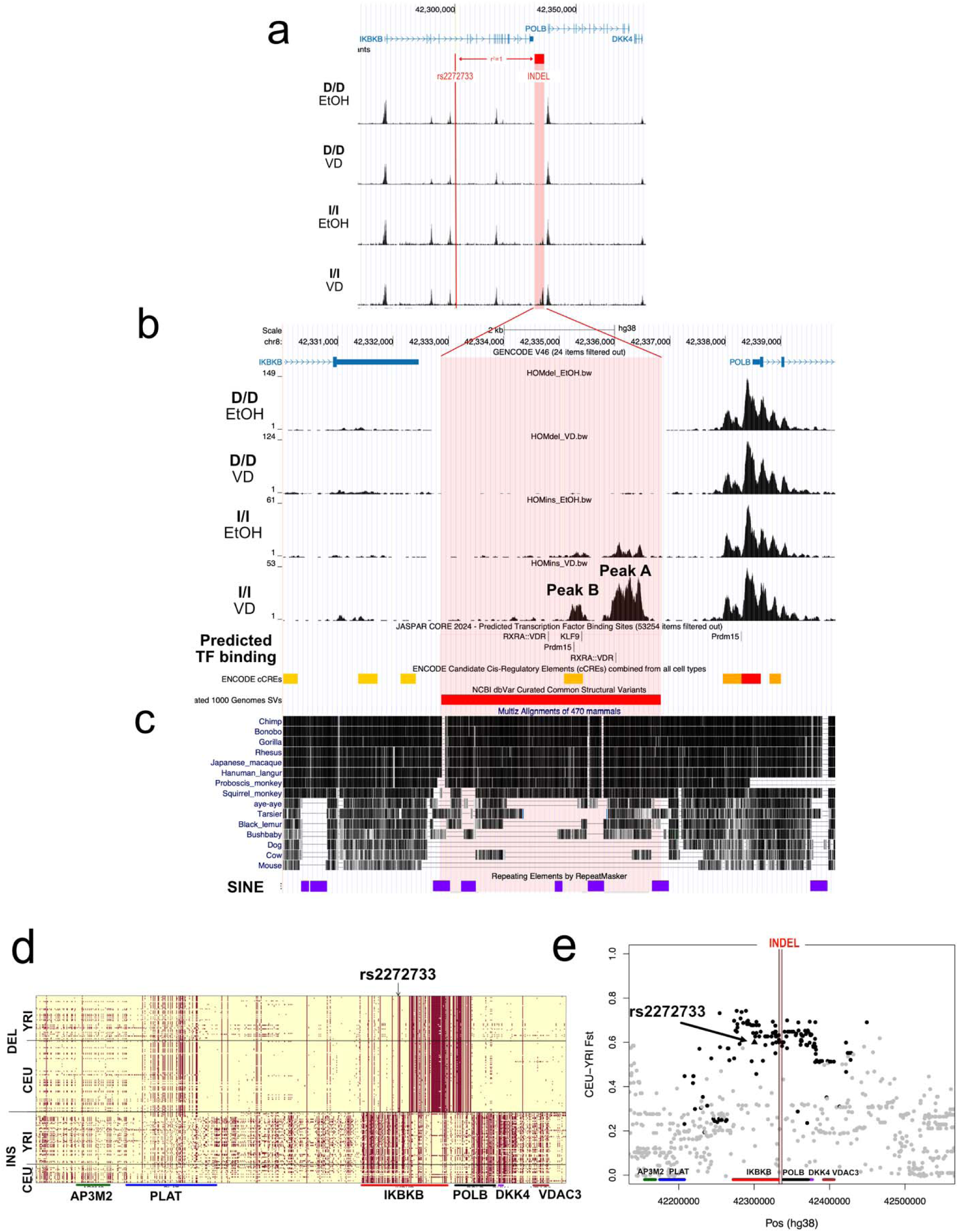
Indel upstream of POLB with large population frequency differences harbors putative regulatory elements and shows signals of natural selection. **4a) rs2272733 is in linkage disequilibrium with an indel.** The SNP rs2272733 is located in an intron of the gene *IKBKB*, while the associated *POLB* proximal peak (“peak A”) is located in a region that overlaps a 4 kb polymorphic 1000 Genome structural indel variant located between the *IKBKB* 3-prime UTR and the *POLB* TSS. The indel was in perfect LD (r^2^=1) with rs2272733 in CEU and YRI. **4b) Insertion harbors two 1,25D responsive peaks with predicted VDREs.** In the insertion, a second, weaker DA peak located 3.1 kb upstream of the *POLB* TSS was identified (“peak B”). Peak B is located in an ENCODE candidate cis-regulatory element with predicted binding of Klf9 and Prdm15. A second strong predicted VDRE is also located in the insertion 3.6 kb upstream of the *POLB* TSS but is not within a peak of chromatin accessibility. **4c) Insertion found to be highly conserved among primates.** The insertion was found to be highly conserved among primates, but not among non-primates. The insertion was flanked by short interspersed nuclear elements (SINEs; shown in purple boxes). **4d) Indel polymorphism individual haplotypes by ancestry.** Comparison of individual haplotypes by ancestry showed haplotype structure more consistent with selection of the derived allele (deletion) outside of Africa. **4e) SNPs with highest F_ST_ associated only with *POLB* responses.** Looking at the highest F_ST_ SNPs in the 200 kb region surrounding the *POLB* indel, we find that they are in high or perfect LD with the indel, significantly associated with *POLB* (black line) expression in 1,25D treatment, and not associated with the expression of any of the nearby genes *AP3M2* (green line), *PLAT* (blue line), *IKBKB* (red line), *DKK4* (purple line), and *VDAC3* (brown line). Combined, these observations suggest that any selection on the indel is likely due to its influence on *POLB* expression. Dot color same as associated egene color; gray dots are not significant eQTLs for any of the shown genes. Triangle represents rs2272733. **Abbreviations:** D/D, homozygous deletion; I/I, homozygous insertion; VD, 1,25D vitamin D; EtOH, ethanol vehicle control; SINE, short interspersed nuclear elements; DEL, deletion; INS, insertion; F_ST_, fixation index; VDRE, vitamin D response element

Taken together, 1,25D responses of *POLB* were of interest due to: 1) ancestry-related transcriptional response differences, 2) a significant vitamin D response *cis-*eQTL, rs2272733, that tagged a haplotype with frequency differences by population, 3) only individuals with the ancestral haplotype showed 1,25D expression and accessibility responses (consistent with a dominant model), 4) *POLB’s* role as a key polymerase in base excision repair^39^ implicated in tumorigenesis,^40-42^ and 5) association of rs2272733 with colorectal cancer^43^. These findings suggest that an ancestral alteration in *POLB* regulation might be an important driver of inter-ethnic differences in responsiveness to 1,25D, with potentially significant consequences for disease risk.

### Indel variant upstream of POLB with large population frequency differences harbors putative regulatory elements and shows signals of natural selection

The core haplotype identified above spans the genes *POLB* and *IKBKB*; however, we found no association with *IKBKB* expression among these SNPs, nor did *IKBKB* show a differential response to vitamin D treatment similar to that seen in *POLB* (**Extended data Figure 4c**). The associated DA peak (“peak A”) is located in a region that overlaps a 4 kb indel located between the *IKBKB* 3’ UTR and the *POLB* TSS (**Figure 4a**), supporting the inference that rs2272733 is a marker for an upstream *POLB* regulatory element. In African and European populations from the 1000 Genomes project (i.e., YRI and CEU; see Methods), this indel was found to be in perfect LD (r^2^=1) with rs2272733 (**Figure 4a**).

In addition to peak A, a second DA peak (“peak B”; LFC=1.56; FDR=5.24x10^-7^) fell within the indel region, 3.1 kb upstream of the *POLB* TSS (**Figure 4b**), signaling a second possible location for our posited vitamin D-responsive regulatory element. Peak A, the stronger, more proximal of the two peaks, harbors a high scoring predicted VDRE (JASPAR score 610) within 250bp of the peak’s center, making it a strong candidate location for the regulatory element we identified. Peak B, the more distal DA peak, is located in an ENCODE candidate cis-regulatory element with predicted binding of KLF9 and PRDM15. Although neither of these two TF motifs is found to be enriched in DA peaks with positive effect (i.e., LFC>0), *PRDM15* expression is significantly upregulated by 1,25D, which could provide a trans-acting regulatory mechanism for this secondary site. There was also a second strong predicted VDRE (JASPAR score 564) located in the insertion 3.6 kb upstream of the *POLB* TSS, although it was not within a peak of chromatin accessibility.

Next, we examined the evolutionary features of the indel to characterize its distribution across ancestral populations and assess whether it has undergone selection based on *POLB* regulation. Mirroring the allele frequency differences for the vitamin D-only eQTL in YRI and CEU populations, in our sample the insertion was present at high frequency in individuals of African ancestry (63%), but less common in those of European ancestry (20%). The insertion is highly conserved among primates, but not among non-primates, and is flanked by short interspersed nuclear elements (SINEs) (**Figure 4c**). To explore the putative age of the indel, we analyzed the AADR ancient DNA database^44^ and noted that the insertion was present in all Neanderthal and Denisovan samples — which is expected given that it is ancestral — but absent in the vast majority of non-African samples older than 5,000 years (**Supplementary Table 9**). These results suggest that the deletion is very old, consistent with the allele having arisen in Africa but having already reached high frequency in Europe more than 15,000 before present (BP).

Given the large allele frequency differences of rs2272733 (and the indel variant in perfect LD), we looked for evidence of differential selection pressure between European and African ancestral regions using data from the 1000 Genomes project. We noted significant test statistics for CEU-YRI F_ST_ (0.586; empirical p=3.0x10^-3^), CEU-YRI cross-population extended haplotype homozygosity (XP-EHH) (2.548; empirical p=1.8x10^-2^), CEU Tajima’s D (-1.913; empirical p=2.0x10^-2^). Taken together, these results are consistent with a positive selection signal within the CEU population. However, we do not find significant statistics for CEU iHS (-0.806; empirical p=2.1x10^-1^), YRI iHS (0.273; empirical p=3.9x10^-1^), or YRI Tajima’s D (-0.932; empirical p=2.0x10^-1^). Comparison of individual haplotypes by ancestry showed haplotype structure more consistent with selection of the deletion outside of Africa (**Figure 4d**), and though neither iHS signal is significant, a negative value in CEU and a positive value in YRI is consistent with selection on the derived allele outside of Africa.

Looking at the highest F_ST_ SNPs in the 200 kb region surrounding the *POLB* indel, we find that they (1) are in high or perfect LD with the indel, (2) are significantly associated with *POLB* expression in 1,25D treatment, and (3) are not associated with the expression of any of the nearby genes *AP3M2*, *PLAT*, *IKBKB*, *DKK4*, and *VDAC3* (**Figure 4e**). Combined, these observations suggest that any selection on the indel is likely due to its influence on *POLB* expression. While the tagging SNP, rs2272733, was not a significant *cis-*eQTL for any gene in colon tissues (sigmoid and transverse colon) in the adult Genotype Tissue Expression (GTEx) project, consistent with our finding of this variant as a condition-specific eQTL, we noted that rs2272733 was a significant *cis-*eQTL for *POLB* in several GTEx tissues (e.g., skin, esophagus, testis, brain, adipose tissue, lung, nerve, skeletal muscle) (**Extended data Figure 5a**). Effects on *POLB* expression in skin, esophagus and, to a lesser extent, testis were in the same direction as effects found in this study (**Extended data Figure 5b**), but opposite in direction in other tissues (**Extended data Figure 5c**). While rs2272733 associations were strongest for *POLB*, other genes, including *RPL5P23, PLAT* and *IKBKB*, were also associated in different tissues in GTEx. Additional studies are needed to understand the role of vitamin D in responses of *POLB* across different tissues to elucidate the observed signals of selection.

**Figure 5:**
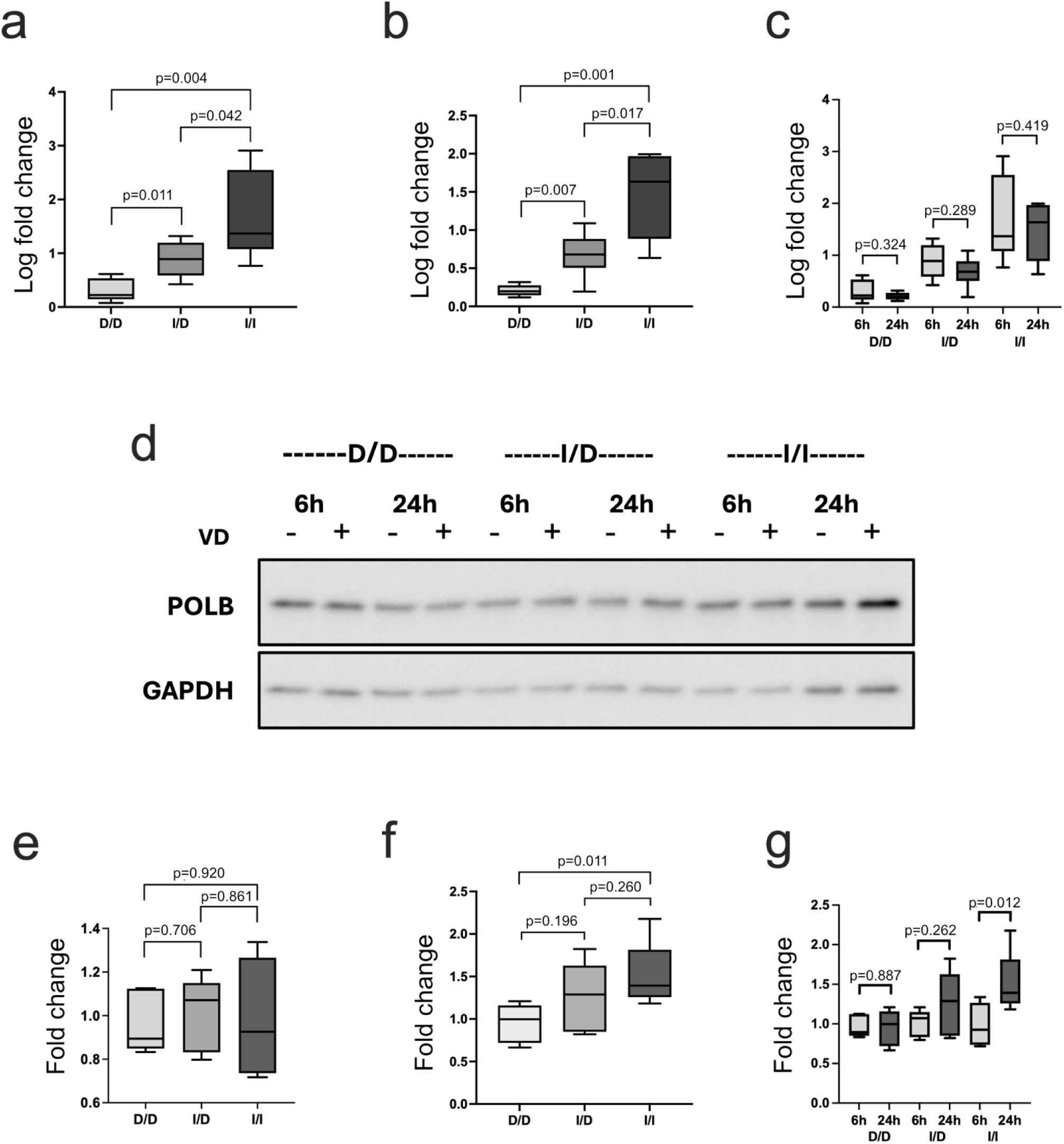
Genotype-specific POLB vitamin D responses and VDR binding. **5a-c) 1,25D responses of *POLB* in organoids at 6 and 24 hours by genotype.** In order to measure *POLB* mRNA expression in response to 1,25D by indel genotype (defined by rs2272733 genotype) at 2 time points, we treated 5 organoid lines representing homozygous deletion (D/D), 6 heterozygotes (I/D) and 7 homozygous insertion (I/I) for 6 (5a) and 24 (5b) hours and measured *POLB* expression by qPCR. Box plots show log fold change expression of *POLB* stimulated by 1,25D in indel genotypes. The levels of expression differed significantly between the 3 groups at the 2 time points tested with the highest, intermediate and lowest seen in I/I I/D and D/D genotypes respectively. No significant changes in *POLB* expression were noted between the time points for all the genotypes (5c). **5d) Western blot of *POLB* protein expression by indel genotype at 6 and 24 hours.** To measure *POLB* protein expression in response to 1,25D by indel genotype (defined by rs2272733 genotype) at 2 time points, we treated D/D, I/D and I/I organoids for 6 and 24 hours and measured *POLB* protein level by Western blotting. Shown are the representative images of blot containing the 3 treated indel genotypes for the 2 time points, probed with antibodies to POLB and GAPDH. Highest level of *POLB* protein was seen in I/I genotype at 24 hours. **5e-g) 1,25D responses of *POLB* protein in organoids at 6 and 24 hours by indel genotype.** For the 6-hour time point, we treated 5 D/D, 5 I/D and 6 I/I organoid lines. For the 24-hour time point, we treated 5 D/D, 5 I/D and 7 I/I organoid lines. Box plots show fold change expression of *POLB* protein by indel genotype for 6 (5e) and 24 (5f) hours. Organoids at 6 hours showed no differences in *POLB* protein levels by indel genotype. The level of *POLB* protein was significantly high in the I/I genotype only at 24 hours. **Abbreviations:** D/D, homozygous deletion; I/D, heterozygote; I/I, homozygous insertion; VD, 1,25D vitamin D

### Genotype-specific POLB vitamin D responses and VDR binding in indel region

To confirm that the *POLB* responses to 1,25D correlated with indel genotype, we identified organoid lines with homozygous deletion (D/D), heterozygotes (I/D) and homozygous insertion (I/I) based on rs2272733 genotype. We performed quantitative PCR (qPCR) to measure *POLB* expression at 6 and 24 hours after 1,25D treatment in D/D (n=5), I/D (n=6) and I/I (n=7). We also used Western blot to measure POLB protein levels in D/D (n=5), I/D (n=5) and I/I (n=7). We observed mRNA induction of *POLB* by 1,25D at both 6 hours and 24 hours (**Figures 5a & b**, respectively). There were significant differences in levels of *POLB* expression between all genotypes. No differences in responses by genotype were noted between 6 and 24 hours (**Figure 5c**).

*POLB* protein expression was not upregulated by 1,25D at 6 hours in any genotype (**Figures 5d & e**). At 24 hours, we noted differential *POLB* upregulation by genotype, with significantly enhanced expression in the treatment group among I/I lines compared to D/D lines and intermediate expression in the I/D lines (**Figure 5d & f**). Only the I/I lines showed significant differences in protein expression between 6 and 24 hours (**Figure 5g**).

To identify the specific elements within the *POLB* insertion that are responsible for the observed genotype-specific 1,25D response, we dissected the transcriptional activity of several regions of interest identified by our ATAC-seq profiling of differential chromatin accessibility. To do this, several vector constructs were designed that included: 1) peak A, the proximal DA peak harboring a predicted VDRE, 2) peak B, the distal DA peak without a predicted VDRE, 3) a second predicted VDRE not in a DA peak, and 4) the region encompassing both peak A and B as well as the two predicted VDREs (**Figure 6a**). We transfected these constructs into 293FT cells, which we exposed to 1,25D or control. After 24 hours, increased transcriptional activity for constructs 1, 3 and 4 was noted, while construct 2 did not show significant transcriptional activity compared to vector control (**Figure 6b**). Notably, the construct that included both chromatin peaks and predicted VDREs showed increased activity compared to the constructs containing each of these elements individually, suggestive of an additive effect of these regions of interest within the insertion.

**Figure 6:**
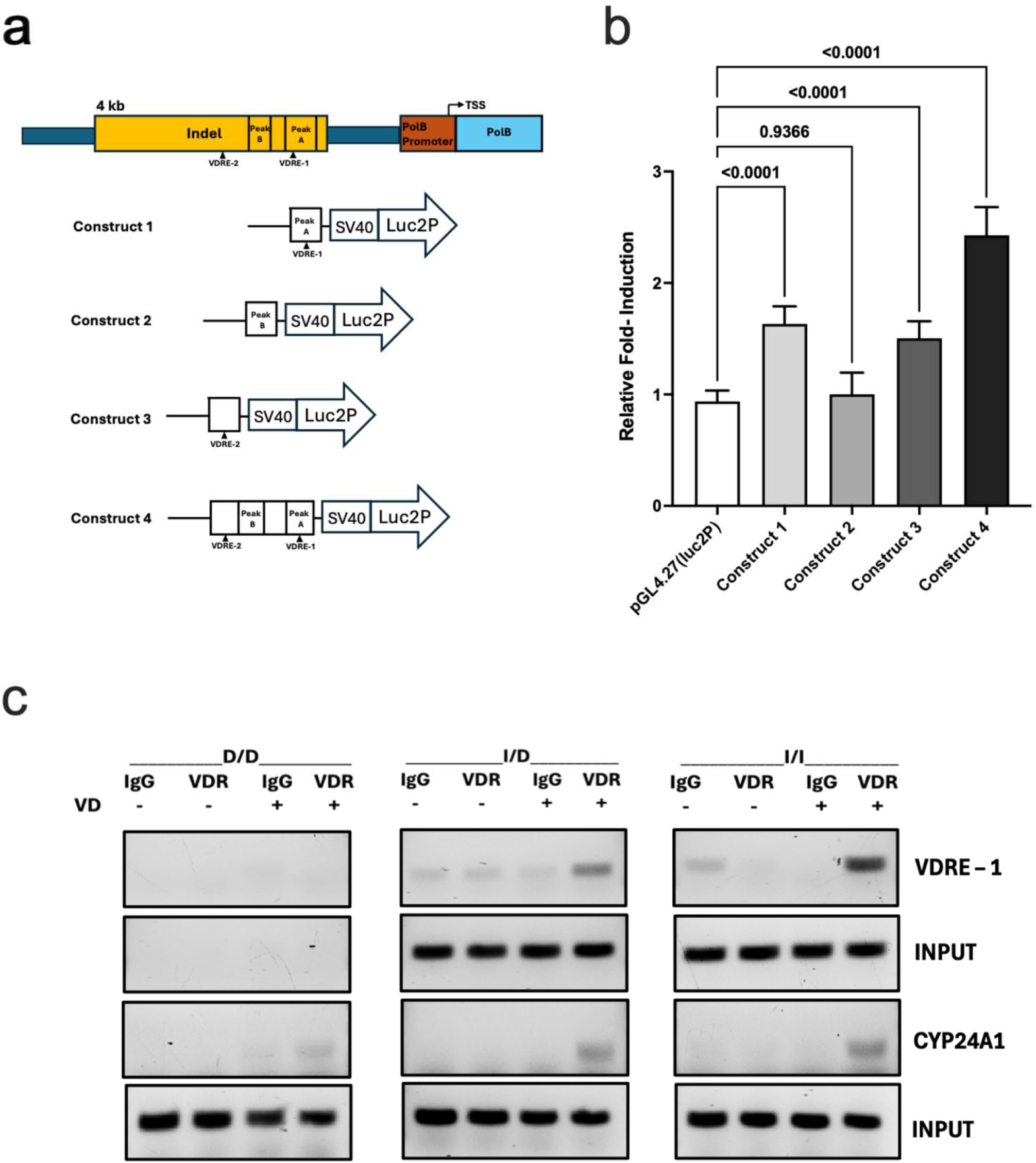
1,25D treatment shows differential transcriptional activity and VDR binding by indel genotype. **6a) Indel region luciferase assay constructs.** Based on results from the ATAC-seq profiling showing differential chromatin accessibility with vitamin D, transcriptional activity of specific regions of interest within the *POLB* insertion was dissected. To do this, several vector constructs were designed that included: 1) the DA chromatin peak harboring a predicted VDRE (“peak A”), 2) the second DA chromatin peak without a predicted VDRE (“peak B”), 3) a second predicted VDRE not in a DA peak, and 4) the region encompassing both peaks A and B as well as the two VDREs. **6b) *POLB* transcriptional activity for different indel constructs.** After 24 hours of 1,25D treatment, increased transcriptional activity for constructs 1, 3 and 4 were noted, while construct 2 did not show significant transcriptional activity compared to vector control. Notably, the construct that included both chromatin peaks and predicted VDREs showed increased activity compared to each construct individually, suggestive of an additive effect of these regions of interest within the insertion. **6c) VDR binding in the indel region by chromatin immunoprecipitation (ChIP) assay.** To provide additional evidence to support differential VDR binding by indel genotype, ChIP assays were performed using organoid lines with homozygous deletion, heterozygous and homozygous insertion. Organoids were treated with or without 1,25D, subjected to ChIP assay, and PCR performed with purified immunoprecipitated chromatin DNA using primers designed to amplify the sequence encompassing the predicted VDRE in peak A. When VDR antibody was added with 1,25D treatment, there was evidence of binding noted in the I/D and I/I lines, while no binding was noted in the D/D line. As a positive control, VDR binding was assessed for a *CYP24A1* promoter region binding site that showed evidence of VDR binding with 1,25D treatment across all genotypes. As a further control, both sequences were amplified from input DNA of all the organoid lines except for the sequence encompassing the predicted VDRE in peak A from the D/D line as expected. **Abbreviations:** D/D, homozygous deletion; I/D, heterozygote; I/I, homozygous insertion; VD, 1,25D vitamin D; VDRE, vitamin D response element; VDR, vitamin D receptor

To provide additional evidence supporting differential VDR binding by indel genotype, chromatin immunoprecipitation (ChIP) assays were performed using organoid lines with homozygous deletion, heterozygous and homozygous insertion. Organoids were treated with 1,25D or vehicle control, subjected to ChIP assay and PCR using primers designed to amplify the sequence encompassing the predicted VDRE in peak A. When VDR antibody was added with 1,25D treatment, there was evidence of binding noted in the heterozygote and homozygous insertion lines, while no binding was noted in the homozygous deletion line **(Figure 6c**). As a positive control, VDR binding was assessed using a proximal human *CYP24A1* promoter region (-252 to -51bp) previously confirmed as a VDR binding site by ChIP-seq in a colorectal cancer cell line.^45^ This region showed evidence of VDR binding with 1,25D treatment across all genotypes. As a further control, both sequences were amplified from input DNA of all the organoid lines except for the sequence encompassing predicted VDRE in peak A from the homozygous deletion line as expected.

## Discussion

Circulating levels of 25D, the inactive form of vitamin D, are known to vary between individuals of diverse ancestries influenced by genetic and environmental factors^46^. In contrast, much less is understood about inter-individual variation in responses to the active form of vitamin D, 1,25D, that mediates a number of important biological functions including protective effects against GI malignant and inflammatory conditions^2-6^. While serum 25D levels are a proxy for vitamin D stores, focus on this measure alone could obscure clinically significant aspects of the 1,25D’s impact on human biology^11^. Our genomic evaluation of 1,25D treatment responses in colonic organoids from individuals of African and European ancestry found a large number of significant alterations in gene expression and chromatin accessibility. We also characterized the role of *cis-*genetic variance on these treatment responses. Integration of results from genomic responses and QTL mapping identified an indel that explains ancestry-associated differences in the vitamin D regulation of *POLB* with signals of positive natural selection. These findings underscore the importance of including samples from genetically diverse individuals in functional genomic studies in order to identify potential drivers of population differences that could be relevant for clinical outcomes and to identify new functionally significant mechanisms that might be obscured by solely focusing on individuals from a single ancestral population.

Our results confirm that short-term 1,25D treatment has broad genomic effects in the colonic epithelium^47^, some of which are genetically regulated. Transcriptional responses to 1,25D were primarily in the direction of gene upregulation and increased chromatin accessibility. DA peaks were enriched in intronic and intergenic regions relative to promoter regions which supports previous reports of vitamin D target gene regulation via super enhancer regions^48^. VDR appeared to be the dominant mediator of genomic effects, and, while several other TFs were found to be enriched among DA peaks, TFs previously reported to co-localize with VDR in a leukemia cell line (i.e., PU.1, CEBPA, GABPA)^49^ were not found to be enriched in the colon. Apart from a handful of candidate genes including *POLB*, our results showed that genomic 1,25D responses were broadly comparable across individuals of African and European ancestry. Our results contrast with large ancestry-specific differences in responses to glucocorticoids^24,50^ and infections^51-53^ across individuals of African and European ancestry in blood and immune cells, which might reflect differences based on selective pressures.

While genetic regulation of circulating levels of 25D has been documented^46^, our finding of significant condition-specific eQTLs across individuals underscores the role of genetic variation in regulation of 1,25D responses, irrespective of 25D levels. Among condition-specific eQTLs, over three-quarters were found under 1,25D treatment. For example, the top eQTL for vitamin D response, rs12672468, was associated with responses of *ZC3HAV1* encoding a zinc-finger protein previously implicated in colorectal cancer^25^ and innate immune responses^26^ that highlights potential new mechanisms of 1,25D actions in the colon that could contribute to differences in disease risks. The only other eQTL mapping study of 1,25D treatment was performed in peripheral blood cells from African Americans and found only a few condition-specific eQTLs^12^, none of which overlapped with the current study, which might reflect differences in tissue responses or experimental approaches.

Integration of results from genomic profiling and QTL mapping identified a novel indel that likely explains observed ancestry-related differences in 1,25D responses of *POLB*, a gene encoding a key polymerase involved in base excision repair. For individuals without at least one insertion (ancestral) allele, *POLB* expression is largely uncoupled from 1,25D exposure, a significant alteration to *POLB* regulation that may be unique in the evolutionary history of primates. Based on our reporter assay and ChIP results, we hypothesize that the regulatory activity occurs via a VDRE located 2.2 kb proximal to the *POLB* promoter within the insertion. A second significant DA peak in the insertion did not have evidence of direct VDR binding but did contain a motif for a different transcription factor, PRDM15, that was significantly upregulated by 1,25D, which suggests the possibility of a trans-acting regulatory mechanism for this second peak. Future work will more precisely define the genetic mechanisms of 1,25D regulation in this region.

The insertion was found to be highly conserved among primates and flanked by short interspersed nuclear elements (SINEs). It is likely that this novel regulatory region was introduced into primates through SINE retrotransposition, specifically Alu-Alu mediated rearrangements^54^. The deletion appears to be present prior to migration out of Africa but rose relatively quickly to higher frequency outside of Africa due to some unknown greater selective pressure. While the exact selective pressures are not known, our results suggest that regulation of *POLB*, rather than other genes in the region such as *IKBKB*, could have been the target for selection, though this hypothesis requires validation in non-colonic tissues. While biological consequences of *POLB* regulation by 1,25D remain to be determined, our findings represent a potential novel mechanism by which 1,25D exerts protective effects against carcinogenesis and, possibly, also inflammation related to oxidative stress in the human colon.

The identification of a vitamin D-dependent regulatory region of *POLB* was particularly interesting given that *POLB* encodes a key enzyme in base excision repair of oxidative DNA damage that is directly involved in colorectal carcinogenesis^40-42^. Studies in both mice^55^ and humans^56^ have shown that vitamin D protects against oxidative stress-induced DNA damage in the colon, and *POLB* could represent a novel mechanism underlying this protective role.

Moreover, the 1,25D-specific eQTL, rs2272733, a tagging SNP for the indel, was previously found to be significantly associated with colorectal cancer; specifically, a protective effect was reported for the T allele^43^ (that tags the insertion). In contrast, in African American head and neck cancer patients, the T allele of rs2272733 was associated with poorer prognosis and treatment responses^57^. We believe that the high F_ST_ and association signals that have been previously reported by others for SNPs in this region are likely driven by their linkage disequilibrium with the indel harboring the *POLB* regulatory element. These contrasting association signals could be explained by pleiotropic tissue- and context-specific effects of vitamin D regulation of *POLB*. Given the central role of *POLB* in DNA base excision repair, even minor perturbations in gene regulation could have significant biological consequences.

In summary, our work leveraging human colonic organoids provides important new insights into mechanisms of context-specific genetic regulation of 1,25D responses in the colonic epithelium. Our results highlight inter-individual differences in responses to the biologically active form of vitamin D on a tissue-specific level, irrespective of serum levels. If confirmed, these findings could inform future efforts to more precisely predict treatment responses for vitamin D-related conditions such as colorectal cancer. We highlight the importance of including diverse individuals in functional genomics research based on identification and mechanistic characterization of a regulatory indel that explains differences in ancestry-related *POLB* responses, findings that broaden understanding of vitamin D regulatory functions with direct implications for health and disease across diverse individuals.

**Extended data Figure 1:**
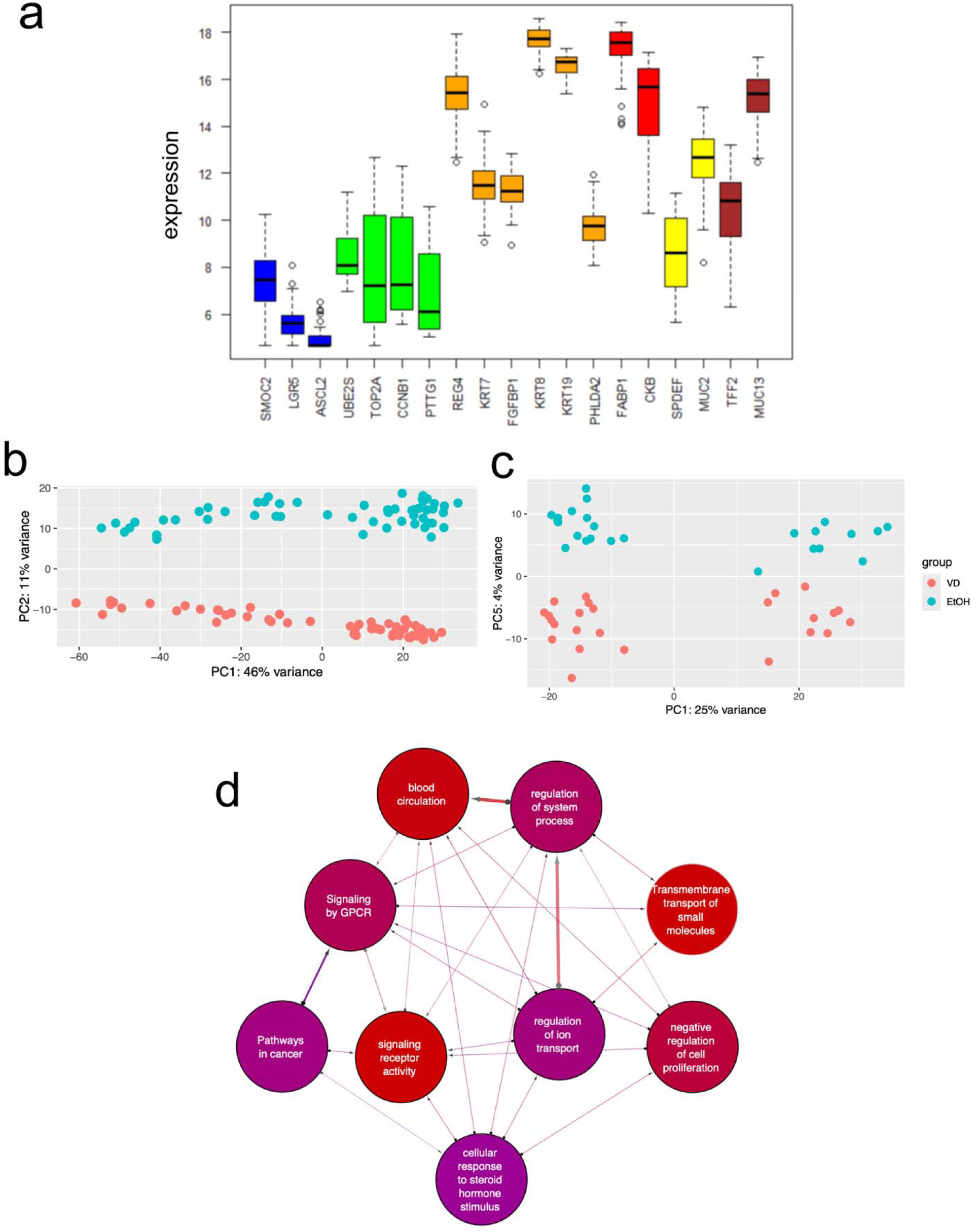
Genomic responses to 1,25D treatment. **1a) Expression of cell type markers in organoids.** Boxplots of the variance stabilized transformed expression levels for canonical cell type markers are shown. The colors are related to cell types: blue, stem cells; green; proliferation stem cells; orange, early enterocytes; red, enterocytes; yellow, MUC2 goblet; brown, MUC13 goblet. Expression is shown for control-treated samples. Expression in control-treated samples was highly correlated with 1,25D-treated samples. No differences in cell type composition was found between populations. To control for confounding due to cell composition, expression of the highly expressed enterocyte markers, *FABP1*, was included as a covariate in the analytical models. **1b) Principal component (PC) plot of transcriptional responses.** 1,25D transcriptional responses measured by RNA-seq were separated by PC2, which accounted for 11% of the variance. **1c) PC plot of chromatin accessibility responses.** 1,25D chromatin accessibility responses measured by ATAC-seq were separated by PC5, which accounted for 4% of the variance. **1d) SetRank network plot.** To visualize interactions of top enriched pathways of DE genes, we used the network output of SetRank plotted using the program Cytoscape (v3.10.2). We show the interactions between the only disease-associated enriched pathway called “pathways in cancer” (KEGG hsa05200) with the top enriched pathways. The node fill color reflects the SetRank corrected p-values with blue to red indicating decreasing p-values. The edge arrows represent interaction from least significant gene set to more significant gene set.

**Extended data Figure 2:**
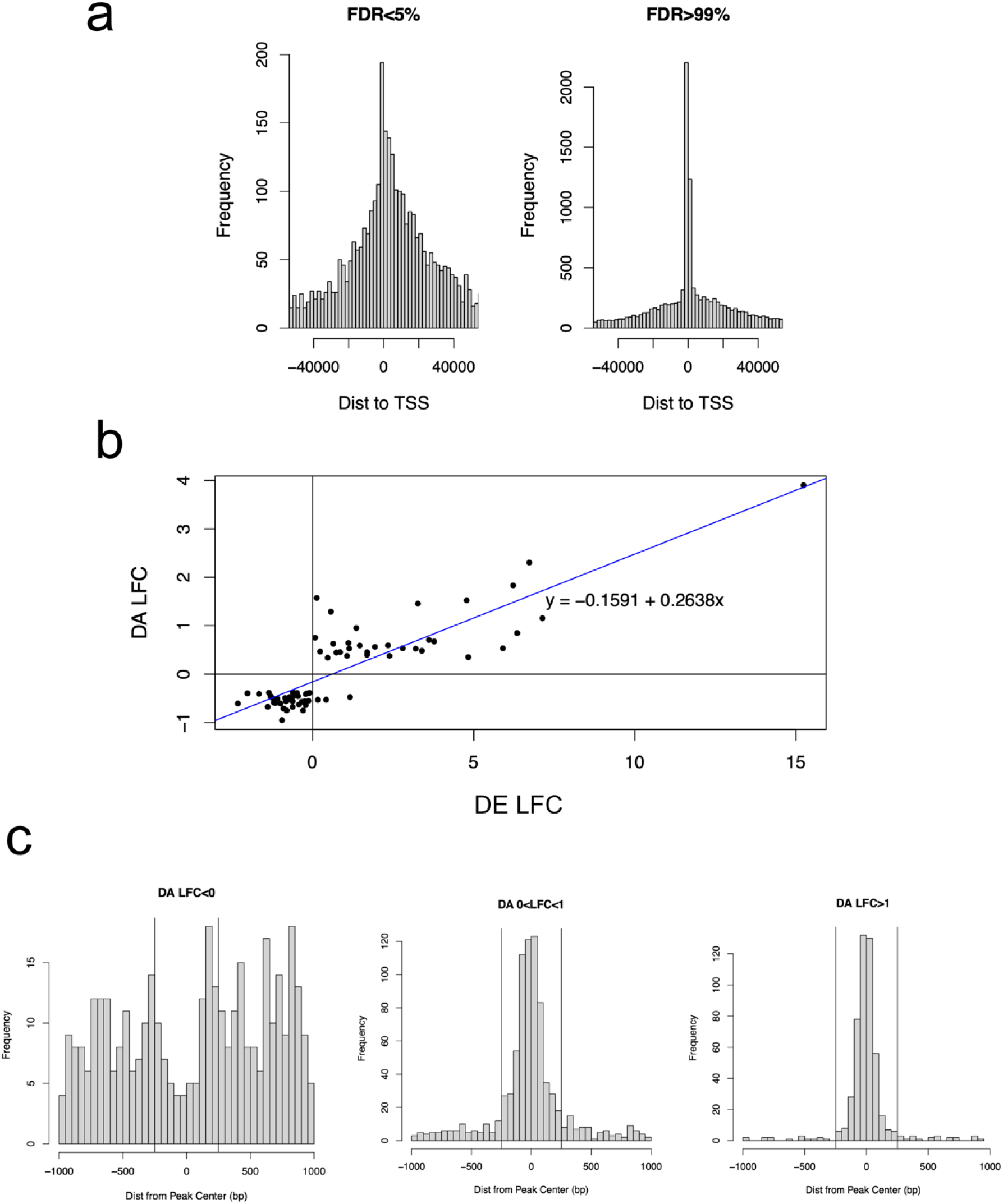
Differentially accessible (DA) peak analysis. **2a) DA peak enrichment.** For DA peaks (FDR<5%), a broader distribution of distances to the transcription start site (TSS) of the nearest gene was observed relative to peaks that were not DA (FDR>99%). This observation is in line with the finding that DA peaks were significantly depleted in promoter regions (3.6% vs. 12.4%, respectively; hypergeometric test p=1.54x10^-89^) compared to intronic or intergenic regions. **2b) Correlation of DE genes and DA peaks in promoter regions.** A total of 85 tested protein-coding genes were found to have DA peaks falling in their promoters. Of these, 73/85 (86%) were DE, a 1.3-fold enrichment over genes without DA peaks in their promoters (hypergeometric test p=8.18x10^-5^). The corresponding DE and DA effect sizes for these 73 genes were strongly correlated (r=0.85; p<2.2x10^-16^) **2c) Vitamin D response element (VDRE) peak enrichment.** The VDRE motif was found within 250 bp of a DA peak center for 75% of DA peaks with large effects (i.e., LFC>1), 46% with moderate effects (i.e.,0<LFC<1), and only 3.8% of DA peaks with reduced effects (i.e., LFC<0). These patterns were similar to VDR enrichment patterns.

**Extended data Figure 3:**
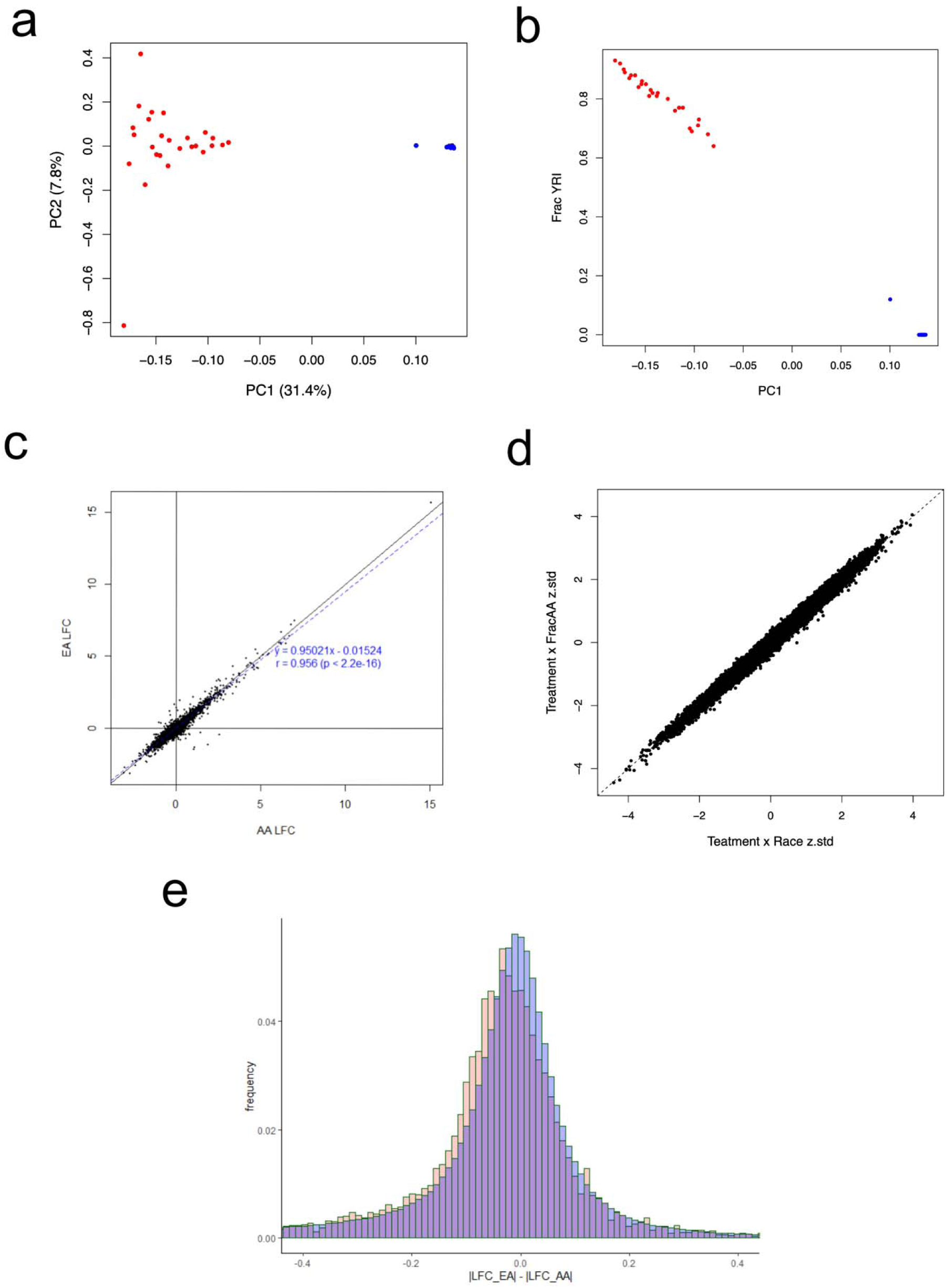
1,25D genomic responses by ancestry. **3a-b) Genetic ancestry.** Genetic ancestry proportions were estimated with the program ADMIXTURE (v1.3.0) using approximately 255,000 imputed SNPs. **a)** Principal component PC1 vs PC2 of genotype SNP data with self-identified White in blue dots and self-identified African-American in red dots, and b) PC1 vs fraction of African (i.e., YRI) ancestry with self-identified White in blue dots and self-identified African-American in red dots. **3c) Correlation of transcriptional response effect sizes in AA and EA lines.** There was significant correlation between effect sizes (i.e., LFC) for DE genes with treatment between AA and EA (r=0.97; p<2.2 x 10^-16^). This result was similar to results for z-scores shown in Fig 1e. **3d) Correlation of treatment effects between models that include fraction African ancestry and self-identified race.** We assessed treatment responses using both fraction African ancestry and self-identified race. There was near perfect correlation between results from the two models (r=0.99; p=2.2 x 10^-16^). **3e) Genome-wide transcriptional responses by ancestry.** To determine whether there were differences in overall absolute response to 1,25D treatment, we compared the between population transcriptional effect size magnitude difference (|LFC_AA_| - |LFC_EA_|) to the expected null distribution generated from permuted samples. Using this approach, neither population showed a stronger overall genome-wide response to treatment (p=0.28).

**Extended data Figure 4:**
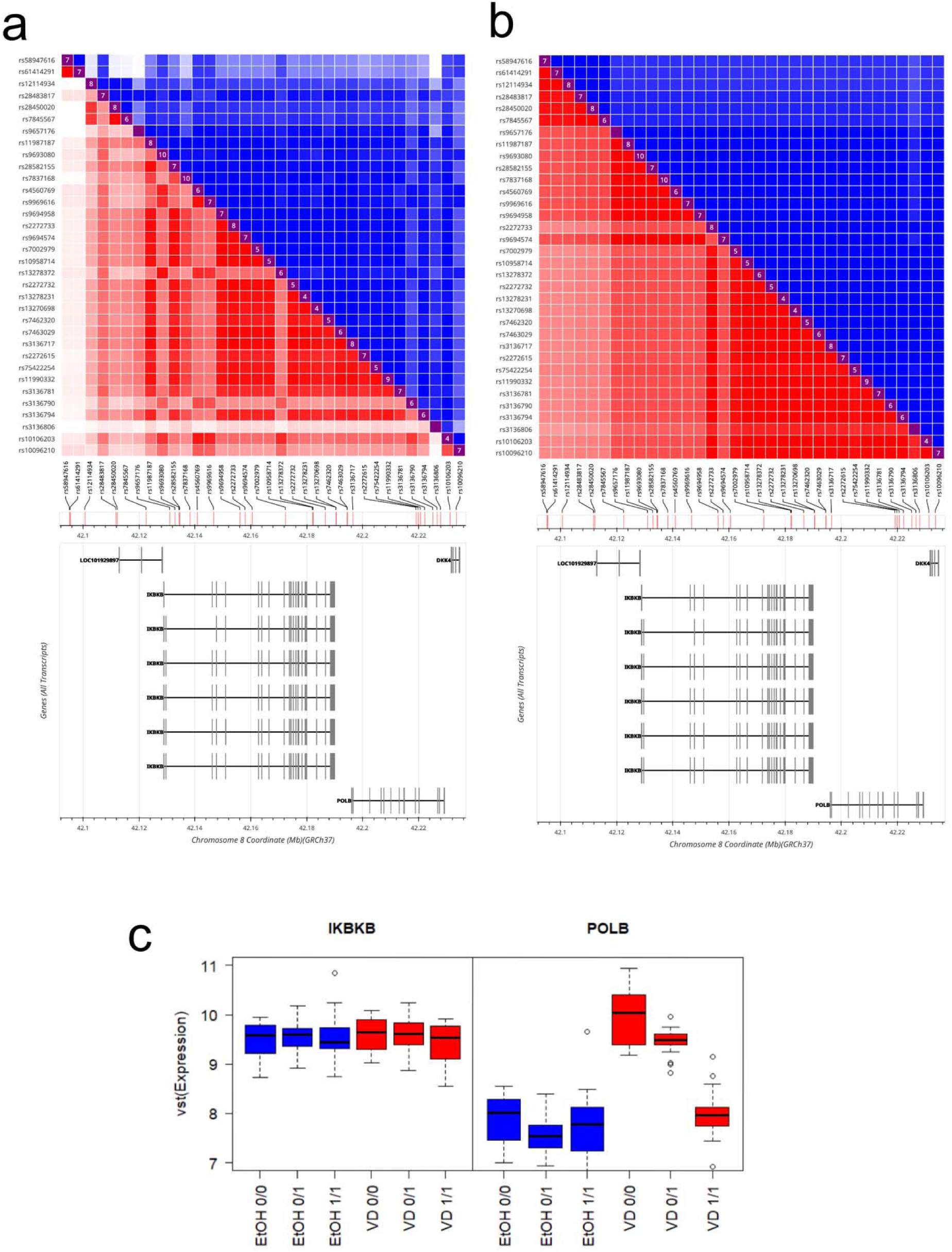
Core haplotypes in *POLB* region. **4a & b) Core haplotype in *POLB* region in YRI and CEU populations.** Of the 34 SNPs that were daQTLs and *POLB* eQTLs, 10 SNPs (rs2272733, rs7002979, rs10958714, rs13278231, rs13270698, rs7462320, rs7463029, rs3136717, rs75422254, rs11990332) were found to define a core haplotype where pairwise SNP r^2^>0.90 in both **4a)** YRI and **4b)** CEU 1000 Genome Project populations. **4c) Core haplotype spans *POLB* and *IKBKB.*** The core haplotype spans the genes *POLB* and *IKBKB*. The tagging SNP rs2272733 showed no association with *IKBKB* expression nor did *IKBKB* show a differential response to 1,25D treatment, while *POLB* showed an association with genotype only in response to 1,25D treatment. Expression shown as the DESeq2 variance stabilized transformation (vst) of normalized read counts.

**Extended data Figure 5:**
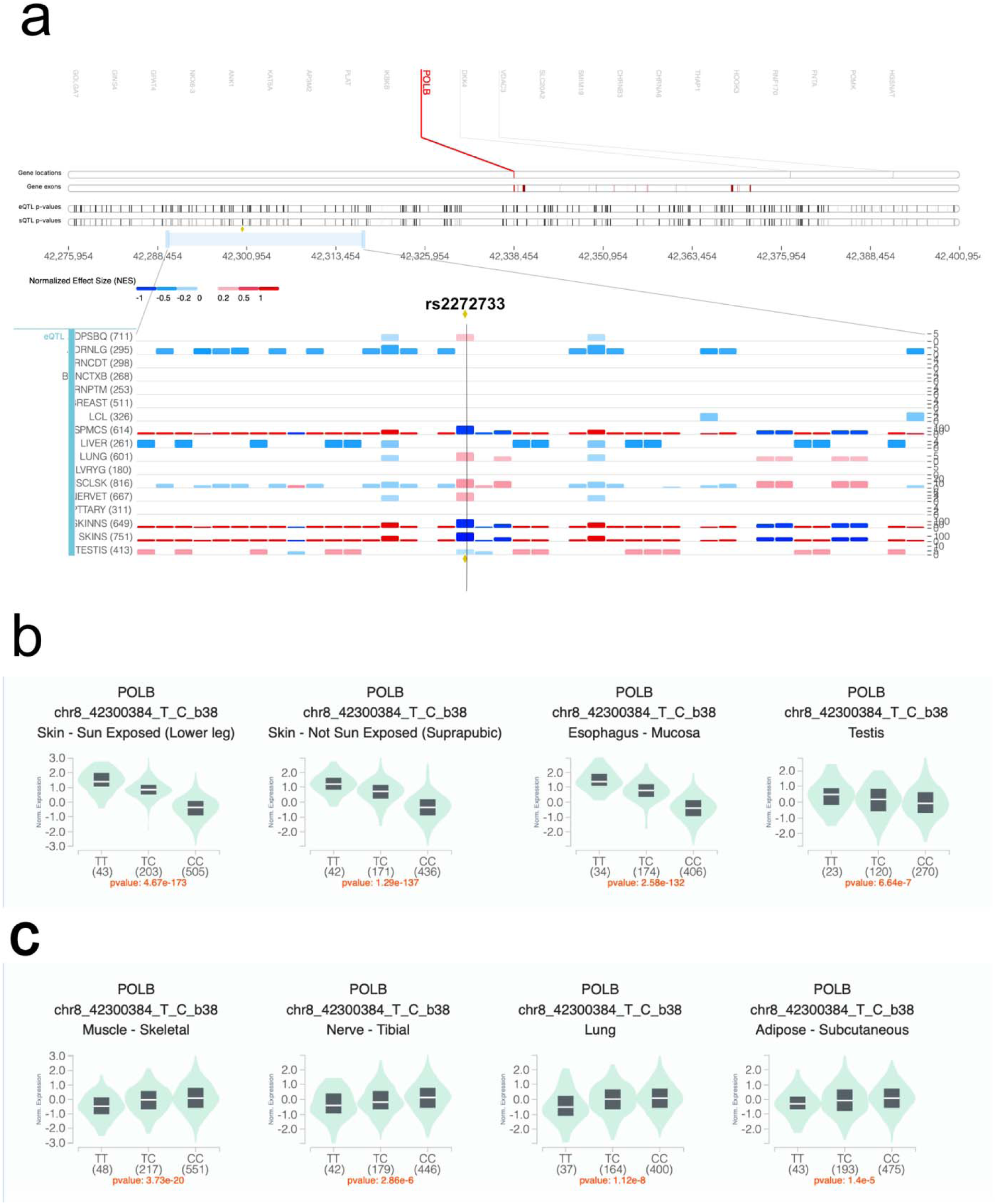
Tissue-specific *cis-*eQTL effects of rs2272733 on *POLB*expression from the Adult Genotype Tissue Expression (GTEx) project. **5a) Effect sizes of rs2272733 in *POLB* expression.** The effects of rs2272733 on *POLB* expression differ by tissue type. Shown here are GTEx tracks for different tissues showing effect sizes by color. rs2272733 is shown as the vertical line. **5b) rs2272733 *cis-*eQTL effects on *POLB* in the same direction as 1,25D colonic responses.** Tissues that showed similar direction of *POLB cis-*eQTL effects for rs2272733 included skin (sun exposed and non-sun exposed), esophagus and, to a lesser extent, testis. **5c) rs2272733 *cis-*eQTL effects on *POLB* in the opposite direction as 1,25D colonic responses.** Tissues that showed opposite direction of *POLB cis-*eQTL effects for rs2272733 included skeletal muscle, nerve, lung and subcutaneous adipose tissue.

## Online Methods

### Study participants

Organoids were derived from rectosigmoid colonic biopsies obtained from consenting participants undergoing screening colonoscopy. Lines from a total of 60 participants were included, comprising equal numbers of women (n=30) and men (n=30) as well as self-identified Black/African Americans (n=30) and non-Hispanic Whites (n=30). The average age of participants was 54.0 years (standard deviation, SD, 9.4) and 57.2 years (SD 9.9) for female and male participants, respectively. The study was approved by the local Institutional Review Board (IRB), and all participants provided informed consent. The final samples included in downstream analyses were determined by quality controls described in more detail in the Bioinformatic section.

### Organoid cultures

Organoids were derived from colonic biopsies using a protocol adapted from Sato et al., as previously described^18,19^. Organoids were cultured on Biolite Multidish (Fisher Scientific, IL), embedded in six 30 µL droplets of Matrigel, cultured in 1.5 mL of the organoid media in an incubator at 37°C and 5% CO_2_. Prior to treatments organoids were incubated in basal media for 24 hours to enable differentiation.

### Treatments

For mRNA expression, organoids were treated with 100 nmol/L 1,25D (Enzo Life Sciences, Farmingdale, NY) or vehicle control (0.1% Ethanol) for 4 hours (for ATAC-seq) and 6 hours (for RNA-seq). For qPCR and Western blotting, organoids were treated for 6 and 24 hours.

### RNA-Sequencing

Organoids were harvested using cold Advanced DMEM, pipetted up and down 10 times, moved to centrifuge tubes, and then spun at 400 g for 5 min at 4°C. Upper media and Basement Membrane Extract (BME) were carefully aspirated and discarded. mRNA was then isolated from cells using the RNeasy® Plus Mini Kit (Qiagen, Germantown, MD) according to the manufacturer’s protocol. RNA quality and quantity were assessed using an Agilent Bio-analyzer with RNA integrity numbers (RIN) of >9 for all samples. RNA-seq libraries were prepared using Illumina mRNA TruSeq Kits as protocolled by Illumina. Library quality and quantity were checked using an Agilent Bio-analyzer, and the pool of libraries was sequenced using an Illumina NovaSeq6000 (paired end 100 bp) using Illumina reagents and protocols at the University of Chicago Genomics Facility.

### ATAC-sequencing

Organoids were harvested at the appropriate time point with collagenase IV and treated with 200 U/mL DNase I (Worthington #LS002007) for 30 minutes. Cells were spun for 5 minutes and DNase 1 supernatant was removed. Cells were then resuspended in cold PBS. The cell density and live cell percentages were measured and then 50,000 live cells per replicate were aliquoted. The cells were then treated using the Omni-ATAC protocol^58^ with some modifications. Briefly, the cells were treated with lysis buffer (10 mM Tris-HCl pH 7.4, 10 mM NaCl, 3 mM MgCl_2_, 0.1% IGEPAL CA-630) and then suspended in the transposition reaction mix (50% TD Tagment DNA buffer, 5% TDE1 Tagment DNA Enzyme, 33% PBS) for 30 minutes. Libraries were prepared using the Nextera Index kit (Illumina #15055289) and were sequenced on the Hi-Seq 4000 platform (50 bp single end reads) at the University of Chicago Genomics Facility.

### Genotyping

Genomic DNA was extracted from blood samples obtained from 64 participants (3 participants did not have available blood samples) and genotyped on the InfiniumOmniExpress-24v1-3_A1 microarray (Illumina; San Diego, CA) which included 714,238 SNPs. Genotype data were used to ascertain genetic ancestry and determine concordance with self-reported ancestry. To increase the density of sites for the purpose of eQTL mapping, genotype imputation was performed with IMPUTE2^59^ using data from 1000 Genomes Project^60^ haplotypes as a reference. Weighted means (dosages) of IMPUTE2’s estimated posterior genotype probabilities were calculated and sites were then filtered by minor allele frequency (>0.05), leaving approximately 8.5 million loci.

### Cell culture

For luciferase assays, 293FT cells were cultured at 37°C in 5% CO2 in DMEM, high glucose (Thermo Fisher Scientific, Waltham, MA) supplemented with 10% FBS and 1% penicillin and streptomycin.

### RNA Isolation, Reverse Transcription and qPCR

After the treatment, RNA was isolated using RNeasy® Plus Mini Kit from Qiagen (Hilden, Germany) as per the manufacturer’s guidelines. Isolated RNA was reverse transcribed into cDNA using a high-capacity cDNA reverse transcription kit from Applied Biosystems, Foster City, CA. qPCR was performed on QuantStudio™ 6 Flex Real-Time PCR System (Applied Biosystems, Foster City, CA) using TaqMan^®^Gene Expression assays (*POLB*: Hs01099715_m1 and *GAPDH*: Hs99999905_m1) from Applied Biosystems, Foster City, CA. Following steps were used for amplification of the target gene: initial denaturation at 95°C for 10 minutes, followed by 40 amplification cycles at 95°C for 15 seconds and 60°C for 1 minute in each cycle. Relative change in expression of *POLB* was analyzed by using the 2^−ΔΔC^_T_ method^61^. Results were expressed as the log fold change in gene expression normalized to endogenous reference gene (*GAPDH*) as well as with the expression of vehicle control at the threshold cycle (Ct). Statistical analysis was performed by means of two-tailed paired t-test with p<0.05 considered as significant.

### Western blotting

Protein was extracted from colonic organoids treated with 100 nmol/L 1,25D or vehicle control (0.1% ethanol) for 6- and 24-hours. Briefly, after treatment, organoids were washed with ice cold PBS and then incubated with RIPA lysis buffer supplemented with protease inhibitor cocktail (Thermo Fisher Scientific, Waltham, MA) for 10 minutes on ice. Lysed organoids were sonicated briefly for 10 seconds, centrifuged at 13,000 rotations per minute for 15 minutes at 4°C and the supernatant was collected. Protein was quantitated using the BCA protein assay kit (Thermo Fisher Scientific, Waltham, MA). Approximately 10-12 µg total protein was used for Western blot assay. The samples were diluted in Laemelli buffer (Bio-Rad, Hercules, CA), boiled for 5 minutes at 95°C. To separate the proteins, samples were subjected to electrophoresis using 4–15% Mini-PROTEAN® TGX Stain-Free™ precast polyacrylamide gels obtained from Bio-Rad, Hercules, CA. Separated proteins were then transferred to a polyvinylidene difluoride membrane. To minimize nonspecific binding, the transferred proteins were blocked using 5% milk in Tris buffered saline with 0.1% Tween 20. The membrane was then probed with primary and secondary antibodies followed by image acquisition on ImageQuant LAS 4000 luminescent imager from (GE Healthcare, Lincoln, NE). Primary antibodies to POLB and GAPDH (catalog numbers ab175197 and 97166S) were obtained from Abcam, Waltham, MA and Cell Signaling Technology Danvers, MA respectively. Secondary antibodies coupled to HRP (catalog numbers 7074S and 7076S) were purchased from Cell Signaling Technology Danvers, MA. Band density was quantitated using ImageJ and normalized to the housekeeping gene to calculate the fold change. Fold change in expression between the vehicle control and 1,25D treated and between the groups was compared using two-tailed paired t-test with p<0.05 considered as significant.

### Luciferase assay

The predicted VDREs in the indel along with a stretch of flanking sequences on their either end was cloned into pGL4.27[luc2P/minP/Hygro] Vector (Promega, Madison, WI) using Infusion^®^ Snap Assembly bundle from Takara Bio USA, San Jose, CA. A stretch of 250 bp sequence without the presence of predicted VDRE was also cloned into pGL4.27[luc2P/minP/Hygro] Vector. DNA was isolated from the positive clones containing the inserted sequences using ZymoPure™ II Plasmid Midiprep kit obtained from Zymo Research Corporation, Irvine, CA, followed by transfection into 293FT cells using Lipofectamine 3000 (Thermo Fisher Scientific, Waltham, MA). Dual-Luciferase(R) Reporter Assay System (Promega, Madison, WI) was used to assess enhancer activity after 24 hours of treatment with 1,25D or vehicle control (0.1% ethanol) according to manufacturer’s guidelines. pRL Renilla Luciferase Control Reporter Vector (pRL-TK) was used as an internal control. Relative enhancer activity was determined by dividing the 1,25D-treated values with ethanol-treated values and compared using one-way ANOVA.

Primers used for generating the constructs:

**Construct 1** (insert size 258 bp with predicted VDRE-1):

Forward: 5’-GAGGATATCAAGATCCCTCTGTTTGGGGAATATTCTATAA-3’ Reverse: 5’-CGCCGAGGCCAGATCGCATTTTAATCCACCCTGCT-3’

**Construct 2** (insert size 250 bp, no VDRE):

Forward: 5’-GAGGATATCAAGATCTATTTCCAGTCCTTCTTAGTACTGT-3’ Reverse: 5’-CGCCGAGGCCAGATCCCACCTCCTGCCTGCTCT-3’

**Construct 3** (insert size 250 bp with predicted VDRE-2):

Forward: 5’-GAGGATATCAAGATCAGCCCATTTCTTGCCCGTAG-3’ Reverse: 5’-CGCCGAGGCCAGATCTGAAGCATGGGACTCTTGGACTC-3’

**Construct 4** (insert size 1480 bp with predicted VDREs-1 and 2): Forward: 5’-GAGGATATCAAGATCCATTTCTTGCCCGTAGCAGTT-3’ Reverse: 5’-CGCCGAGGCCAGATCGCATTTTAATCCACCCTGC-3’

### Chromatin immunoprecipitation (ChIP)

ChIP assay was performed using the SimpleChIP® Plus Sonication Chromatin IP Kit from Cell Signaling Technology, Inc. (Danvers, MA) following the manufacturer’s instructions. Briefly, organoids were treated with EtOH or 100 nM 1,25D for 6 h, dissociated into single cells by TrypLE Express and cross-linked using 1% methanol free formaldehyde. Cells were then lysed, and the chromatin pellets were fragmented by sonication to an average size of 200- to 1000-bp, using a Fisherbrand Model 120 Sonic Dismembrator (Thermo Fisher Scientific, Waltham, MA). Precleared sonicated extract was diluted into ChIP buffer and subjected to immunoprecipitation by incubating with either a control IgG antibody or 2-4 μg of mouse monoclonal antibody to VDR (Santa Cruz Biotechnology, Dallas, TX) overnight on a rotator at 4 C. The immunoprecipitated DNA fragments were eluted, purified and subjected to PCR using the following pair of primers for predicted VDRE in the indel: forward, 5’-GACAAGAGCAGAAGCAGGAA-3’; reverse, 5’-CTATCAGGCCAAACCCATAAGA-3’, which were designed to amplify the indel sequence from coordinates chr8:42193437 to – chr8:42193648 encompassing predicted VDRE and give rise to a 212-bp fragment. A sequence 241 bp encompassing VDRE in the *CYP24A1* promoter was amplified using the primers: forward, 5’-CGAAGCACACCCGGTGAACT-3’; reverse, 5’-CCAATGAGCACGCAGAGGAG-3’ from Meyer et al^62^ and used as a control. PCR products were resolved on 1% agarose gels and visualized using Sybr safe staining. DNA acquired before precipitation was used to assess the presence of sequence to be amplified following the ChIP procedure and designated “Input.”

## Bioinformatic analyses

### Ancestry estimates

Genetic ancestry proportions were estimated with the program ADMIXTURE^63^ (v1.3.0) using approximately 255,000 imputed SNPs that were pruned for linkage disequilibrium and filtered for a minor allele frequency greater than 0.05 using PLINK2^64^ (v2.00) commands –indep-pairwise 50kb 10 0.2 and –maf 0.05. The genotype data were then merged with the genotype data from 10 CEU, 10 YRI, and 10 CHB from the 1000 Genome Project. ADMIXTURE was run with k=3 to capture African, European, and possibly Native American or Asian ancestry components of individuals who self-identified as either non-Hispanic White or Black/African American. Principal component analysis (PCA) was performed on the same set of SNPs using PLINK2 with the --pca command.

### Transcriptional responses

Sequence alignment and gene expression value estimation was performed using the rsem-calculate-expression function of the RSEM^65^ v.1.3.1 software package using the STAR^66^ aligner. The STAR transcriptome reference was generated from the 1000 genomes Phase2 Reference Genome Sequence (hs37d5) and transcript annotations from the Gencode comprehensive gene annotation GTF (Release 29). As QC measures all mapped RNA-seq samples were checked for both pairwise relatedness and ancestry proportions. The programs angsd^67^ (v0.941-17-ge6967e6) and ngsRelate^68^ (v2) were used on the resulting alignment files to first estimate genotype likelihoods and then pairwise sample relatedness.

Using the angsd-generated genotype likelihoods, ancestry admixture proportions were estimated with the program fastNGSadmix^69^ and the CEU, YRI and CHB data from the fastNGSadmix 1000 Genomes reference panel. Individuals were removed from downstream analyses if any of the following were found to be true: (1) the individual’s VD and EtOH treatment samples were genetically unrelated; (2) either treatment sample was genetically related to a sample from a different individual; (3) either treatment sample showed an ancestry proportion dissimilar to that estimated from the genotype data; and (4) expression of the canonical vitamin D-responsive gene, *CYP24A1*, was discordant with treatment labels.

Genes tested for differential expression were initially filtered for protein coding biotype and expression level by applying a minimum total count threshold of 10 across all samples and then using the filterByExpr function from edgeR^70^ (v4.0.16) R package (all R packages were run in R v4.3.1).To account for the paired nature of the data (2 treatment samples per individual) and the inclusion of additional covariates, differential expression was tested with mixed linear models using the dream statistical package^20^, which is part of the variance Partition R package^71^ (v 1.32.5) and is built on top of the standard limma^72^ (v 3.58.1) workflow. In all models, the individual term was treated as a random effect, while covariates and predictor variables were treated as fixed effects. Model covariates included Batch, Age, and Sex ascertained from genotype data. Single cell analysis of an organoid line after 24 hours in differentiation media was performed. At this time point, cell populations included stem cells, proliferating stem cells, early enterocytes, enterocytes and goblet cells (**Extended data Figure 1a**). *FABP1*, a known marker for enterocytes, was found to be the most highly expressed cell-specific gene and so expression of this gene was included as an additional covariate in the models to control for potential confounding by cell type composition among organoids. Of note, no differences in cell type markers was observed by ancestry. The following mixed effects Model 1 was used to test for overall differential expression in response to treatment averaged over both populations:

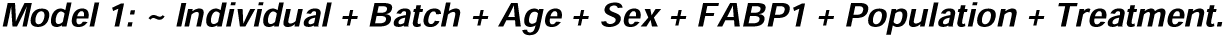

To test for differences in response to treatment within each population, differences of populations within each treatment and differences of response to treatment between populations, an interaction term whose coefficient represents the difference in response to treatment between the EA and AA populations was added to Model 1 to create Model 2a:

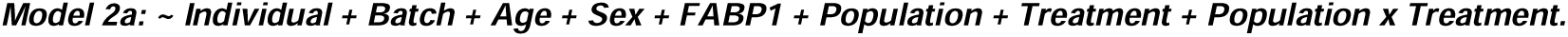

In all cases, the false discovery rate (FDR) was controlled using a Benjamini-Hochberg (BH) adjustment of the estimated p-value. An FDR of 5% or less was considered significant unless otherwise indicated. Because the power for detecting significance is reduced in the second order interaction term in Model 2a, the set of genes tested for this term was restricted to those that showed both DE significance by response to treatment in either EA or AA population and DE significance by population in either treatment condition. To test whether genetic ancestry and self-reported ancestry yielded similar results, the following Model 2b was also applied:

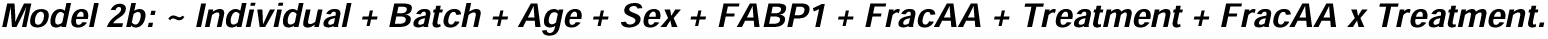

where FracAA is the proportion of African ancestry as reported by ADMIXTURE.

In box plots, gene expression is reported as the transformed and normalized counts after applying the variance stabilizing transformation (VST) of DESeq2^73^ (v1.44.0). Individual differential gene expression is computed from these VST values and reported as the log_2_ fold change of VD/EtOH.

### Gene set enrichment analysis

Gene set enrichment analysis (GSEA) was performed using the R package SetRank^21^. This method was designed to minimize false positives by taking gene set overlap into account. Briefly, the algorithm inputs a gene set collection from the KEGG, GOBP and REACTOME databases and the list of differentially expressed genes in response to treatment (not filtered by a p-value cut-off). The output includes a setRank value (reflects the importance of the gene set in the gene set network; the higher the value, the more important the gene set), a p-value associated with the SetRank value (probability of observing a gene set with the same SetRank value in a random network), a corrected p-value (account for overlap with other gene sets) and adjusted p-value (correction of multiple testing). Pathways with the highest SetRank values and associated SetRank p-values<0.05 are shown in the results. The gene set network was created from the SetRank output using the software Cytoscape^74^ (v3.10.2). The gene set nodes selected for display were the single significant disease-related gene set (KEGG: Pathways in cancer) and all pSetRank significant nodes.

### Chromatin differential accessibility analysis

ATAC-seq read alignment was performed with bwa mem (bwa 0.7.17). Reads were mapped both to the 1000genomes Phase2 Reference Genome Sequence (hs37d5) and the Homo_sapiens_assembly38.fasta assembly (downloaded from the UCSC Genome Browser website) after the removal of alternate haplotypes. All ATAC results are given for the build 37 assembly. Read filtering was performed with Samtools^75^ (v1.10) retaining only uniquely mapped reads with a mapping quality >=10. PCR duplicates were removed with the samtools markdup function. Similar to the RNA-seq samples, ATAC samples were checked for sample relatedness and ancestry proportion using the ngsRelate and fastNGSadmix tools, respectively, and individuals were removed from the ATAC differential accessibility analysis if any of the following applied: (1) the individual’s VD and EtOH treatment samples were genetically unrelated; (2) either treatment sample was genetically related to a sample from a different individual; or (3) either treatment sample showed an ancestry proportion dissimilar to that estimated from the genotype data. On the retained samples, ATAC peak calling was performed with MACS2^76^ (2.1.0) using the callpeaks function with the arguments --nomodel, shift=-100, and extsize=200. The following three criteria then had to be met for individual inclusion in the downstream ATAC DA analysis:

(4) MACS peak count was >30,000 for both treatment samples; (5) fraction of reads in peaks (FRiP) > 10% for both treatment samples; and (6) a significant DA peak in the CYP24A1 promoter with LFC(VD/EtOH)>1. After applying these quality filters, ATAC-seq data from 25 organoid lines were included (10AA and 15EA; 15 females and 10 males). In samples mapped to hs37d5, MACS2 called between 30,130 and 111,962 peaks at an FDR less than or equal to 5%, and FRiP scores were between 11% and 36% with an average of 22.4%. Peak differential accessibility analysis was performed with the R package DiffBind^22^ (v3.12.0) by employing edgeR (v4.0.16) as the underlying method for the differential peak read count analysis. For the combined population analysis, the DiffBind pipeline was run on a consensus peakset formed from the set of MACS2 peaks present in at least 3 of the 50 samples across both treatments (118,806 peaks). Count data normalization by library size and a standardized differential analysis were performed with edgeR. edgeR normalization factors were computed using the TMM method without precision weights, and then the GLM pipeline was run with tagwise dispersion estimates. Differential binding in response to treatment was tested across the 118,806 peakset in both the entire sample set and the separate AA and EA populations using the following Model 3:

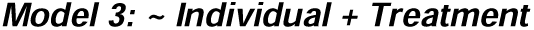

A likelihood ratio test was performed for hypothesis testing and FDR controlled using a BH adjustment of the estimated p-value.

Motif enrichment and peak annotation was performed with HOMER^23^ (v5.1) using the built-in set of HOMER transcription factor motifs. For HOMER analysis of the set of differentially accessible peaks, the background was chosen to be the set of all 118,806 peaks analyzed for differential accessibility. FDR of 5% or less was considered significant for motif enrichment calls.

### cis-eQTL analyses

*Cis*-eQTL mapping was performed on the expression data of protein coding genes after adjusting for batch effects with ComBat-seq^77^ (part of the sva v3.52.0 R package), VST transformation with DESeq2, and quantile normalization, and using imputed bi-allelic variants that fall within 500 kb of a gene’s transcription start site and have a MAF>0.05. After applying these filters, there were approximately 105.5 million gene-variant pairs for analysis. To find condition-specific eQTLs, the program BRIdGE^24^ was run, using the documentation suggested grid of effect sizes. A posterior probability threshold of 0.70 was used to assign significance to an eQTL that had either an VD effect only, an EtOH effect only, equal effects in both treatment conditions, or unequal, nonzero effects in both conditions. Additionally, cis-eQTL mapping was performed separately on the VD and EtOH normalized expression with the R package MatrixEQTL^37^ (v2.3) In VD, 6,908,303 SNP-gene tests had nominal p-value<0.05 with 39,804 having FDR<0.05; in EtOH, 6,518,536 SNP-gene tests had nominal p-value<0.05 with 32,487 having FDR<0.05. Results from MatrixEQTL showed good concordance with those from BRIdGE. We also found that 49% of *cis*-eQTLS identified by BRIdGE overlapped with eQTLs reported in the Adult Genotype-Tissue Expression project (GTEx v7) dataset across 48 tissues.

To explore potential functional relevance of these condition-specific eQTLs for human traits and diseases, we assessed overlap of these variants (or those in strong linkage disequilibrium) with variants from several sources: 1) publicly available databases (e.g., UK Biobank and GWAS catalog) and 2) previously published studies.

### Chromatin differential accessibility QTL analyses

cis-daQTL mapping was performed on the LFC of significant differentially accessible ATAC peaks for the 25 individuals with passing ATAC QC metrics, using all imputed bi-allelic SNPs falling within 100 kb of peak start or end positions. Peak LFC was calculated from the DESeq2 VST transformation of peak read counts, and association mapping was performed using the program MatrixEQTL (v2.3), applying both linear and ANOVA models to detect additive and dominant associations, respectively. For the linear model, of 2,110,074 DA peak-SNP tests, 132,578 tests had nominal p-value<0.05 with 594 having an FDR<0.05; for the ANOVA model, of 2,110,074 tests, 106,277 tests had nominal p-value<0.05 with 2,108 having an FDR<0.05.

### POLB selection signal analyses

iHS values reported for rs2272733 in 1KG Phase 3 CEU and YRI populations based on the approach from Johnson KE et al^78^. applying their updated normalization method; associated empirical p-values are estimated from the normal distribution function. Weir Fst, cross-population extended haplotype homozygosity (XP-EHH) and Tajima’s D statistics are from Pybus M et al^79^; associated p-values are calculated from genome-wide rank scores. The visual haplotype was created from 1000 Genome Project phased genotype data for CEU and YRI.

### Inference of POLB INDEL genotype

The genotype of the INDEL (nssv16196380) proximal to the POLB promoter is imputed in our samples using the 1000 Genome Project Phase 3 dataset as a reference. We also more directly infer INDEL genotype from the genotyped SNP rs2272733 because the two are in perfect LD (r^2^=1) in the 1KG CEU and YRI samples, as well as in our samples. We also validated genotype status of samples inferred to be homozygous deletion from ATAC-seq coverage of the region.

